# Regulation of lipolysis by 14-3-3 proteins on human adipocyte lipid droplets

**DOI:** 10.1101/2023.05.01.538914

**Authors:** Qin Yang, Zinger Yang Loureiro, Anand Desai, Tiffany DeSouza, Kaida Li, Hui Wang, Sarah M Nicoloro, Javier Solivan-Rivera, Silvia Corvera

## Abstract

Adipocyte lipid droplets (LDs) play a crucial role in systemic lipid metabolism by storing and releasing lipids to meet the organism’s energy needs. Hormonal signals such as catecholamines and insulin act on adipocyte LDs, and impaired responsiveness to these signals can lead to uncontrolled lipolysis, lipotoxicity, and a higher risk of metabolic diseases. To investigate the mechanisms that control LD function in human adipocytes, we employed techniques to obtain mesenchymal progenitor cells on a large scale and applied proximity labeling mediated by enhanced ascorbate peroxidase (APEX2) to identify the interactome of PLIN1 in differentiated adipocytes. We identified 70 proteins that interact specifically with PLIN1, including PNPLA2 and LIPE, which are the primary effectors of regulated triglyceride hydrolysis, and four members of the 14-3-3 protein family (YWHAB, YWHAE, YWHAZ, and YWHAG), which are known to regulate diverse signaling pathways. Functional studies showed that YWHAB is required for maximum cAMP-stimulated lipolysis and helps to mitigate the anti-lipolytic effects of insulin. These findings reveal new regulatory mechanisms that control lipolysis in human metabolism.

**SIGNIFICANCE STATEMENT:** Lipid droplets are ubiquitous cytoplasmic organelles that store metabolic energy and play a key role in cellular lipid metabolism (1). Adipocyte LDs play an additional, crucial role, as they supply the energy needs of the whole body through hormonally regulated triglyceride synthesis, storage, and release. The mechanisms by which adipocyte lipid droplets release lipids for systemic use has been mostly studied in mouse models and cell lines. To understand how lipid mobilization is controlled in human adipocytes, we used proximity labeling to identify proteins that interact with PLIN1, a major component of the lipid droplet, in adipocytes generated from primary human progenitor cells. Our study catalogues the interactome of human PLIN1 and identifies previously unrecognized potential mechanism for control of human adipocyte lipolysis through 14-3-3 proteins.

## INTRODUCTION

Lipid droplets (LDs) are unique organelles that store neutral lipids, have a ubiquitous presence in eukaryotic cells, and are considered central hubs for cellular lipid metabolism, signal transduction and trafficking events(1–3). In adipocytes, LDs play the additional key role of providing for the storage and supply needs of the whole organism, a function that is controlled largely through catecholamine and insulin signaling. Under fed conditions, insulin inhibits lipolysis and promotes esterification of fatty acids into triglycerides, which are stored in the large lipid droplets characteristic of differentiated adipocytes. The failure to store lipids appropriately in adipocytes can lead to excessive lipid accumulation in cells of other tissues such as liver, muscle, and heart, leading to lipotoxic oxidative stress and mitochondrial damage during autophagy (4). Under conditions of starvation, exercise or cold-exposure, neutral lipids stored in LDs are rapidly mobilized and enter the circulation to provide for the energy needs of other tissues. Given the central role of adipocyte LDs in systemic energy homeostasis, dysregulation or dysfunction of LDs has major impact on human metabolic diseases such as obesity, diabetes, cardiovascular disease, and non-alcoholic fatty liver disease.

In mammalian cells, LDs consist of a hydrophobic core of neutral lipids such as sterol esters and triacylglycerols, surrounded by a phospholipid monolayer decorated by numerous specific proteins (2, 3). During fed conditions, fatty acids are esterified into triglycerides in the endoplasmic reticulum and directed to lipid droplets for storage by mechanisms that are controlled by insulin. Under fasting, exercise or cold exposure, catecholamine signaling stimulates hydrolysis of LD triglycerides into fatty acids and glycerol (5) in a process mediated sequentially by the three lipolytic enzymes adipose triglyceride lipase (ATGL, *PNPLA2*) (6), hormone-sensitive lipase (HSL, *LIPE*) (7) and monoacylglycerol lipase (MGL, *MGLL*) (7). The regulation of fatty acid esterification and triglyceride lipolysis is mediated by proteins that interact with the surface of the LD. Under basal conditions, perilipin 1 (*PLIN1*), the most abundant protein localized on the surface of LDs (8–10), sequesters CGI-58, a co-activator of ATGL, preventing lipase activity (11). In response to catecholamines, PLIN1 becomes phosphorylated by protein kinase A (PKA), releasing CGI-58 which can then activate ATGL to initiate lipolysis (11). HSL is also phosphorylated by PKA and then translocated from the cytosol to the surface of LDs, thereby exerting the hydrolytic enzyme activity.

This most widely accepted model of lipolysis regulation is mainly derived from murine cells and mouse genetic models, but whether identical mechanisms operate in human adipocytes is largely unknown. There are important differences between mouse and human adipocytes, including significant differences in cell and lipid droplet size which could influence regulatory mechanisms. Indeed, in a previous study, we found that a primate-specific long noncoding RNA, *LINC00473* colocalizes with PLIN1 and positively regulates stimulated lipolysis(12), but an analogous molecule has not been identified in mice. Detailed understanding of LD proteome dynamics triggered by lipolytic stimuli can ultimately help us to reveal the human-specific mechanisms controlling lipid mobilization in a more comprehensive way.

The identification of LD proteins has predominantly been accomplished using subcellular fractionation combined with proteomic analyses, using mouse clonal cell lines (13, 14) or mouse adipose tissues(15, 16). More recently, proximity labeling by APEX2 (17, 18) has been developed as a powerful technique to identify the interactome of specific proteins in intact cells. Using H_2_O_2_ as a co-substrate, APEX2 rapidly oxidizes biotin-phenol (BP) into a highly reactive biotin-phenoxyl radical, which diffuses outward to covalently tag proximal endogenous proteins within 10 nm with fast kinetics (< 1 min) and high activity. PLIN2-APEX2 and ATGL(S47A)-APEX2 chimeras were used recently to characterize the composition of the LD proteome in two different cell lines, U2OS and Huh7 (19). However, these cells do not exhibit the large, hormonally responsive LDs characteristics of adipocytes. Here, we used a similar approach to identify the PLIN1 interactome in human adipocyte lipid droplets under different conditions of hormonal stimulation. We leveraged techniques to obtain human adipose tissue-derived progenitor cells at scale (18), which have the capacity to differentiate into at least four adipocyte subtypes (20). Our studies identify additional members of the LD proteome, including four members of the 14-3-3 family (YWHAB, YWHAE, YWHAG, YWAHZ). Functional studies reveal that YWHAB modulates cAMP-induced stimulated lipolysis and the anti-lipolytic action of insulin.

## RESULTS

### Generation and characterization of APEX2 fusion constructs

Three lentiviral vectors were constructed: a V5-tagged APEX2 genetically fused to the C-terminus of PLIN1 (PLIN1-APEX2-V5), a cytosolic version of APEX2 as a spatial reference control (Cyto-APEX2-V5), and an empty vector served as a negative control (Fig. 1A). Cells transduced with each construct were stimulated with forskolin (Fsk) for 6h to induce lipolysis, and then incubated with biotin-phenol for 30 min. H_2_O_2_ was added for 1 min to initiate labeling, after which cells were fixed for immunofluorescence analysis or lysed in a boiling buffer containing 2% SDS. Biotinylated proteins in the diluted lysates were recovered using streptavidin or neutravidin beads for further identification by LC-MS/MS (Fig. 1B). We first determined the subcellular localization of the APEX2 constructs by immunofluorescence using antibodies against V5 tag (Fig. 1C). Cyto-APEX2-V5 was diffusely distributed throughout the cell including cytoplasm and nucleoplasm. By contrast, PLIN1-APEX2-V5 was localized to the periphery of LDs, which were visualized by LipidTOX staining. To determine whether protein biotinylation would be restricted to the vicinity of the expressed constructs, we visualized the biotinylated proteins using streptavidin Alexa Fluro-568. A diffuse, whole-cell labeling pattern was observed in cells expressing Cyto-APEX2-V5, but biotinylation in cells expressing PLIN1-APEX2-V5 was observed on the periphery of LDs, either in the absence or presence of Fsk stimulation (Fig. 1D), confirming that labeling was restricted to the vicinity of each construct.

**Figure 1.**
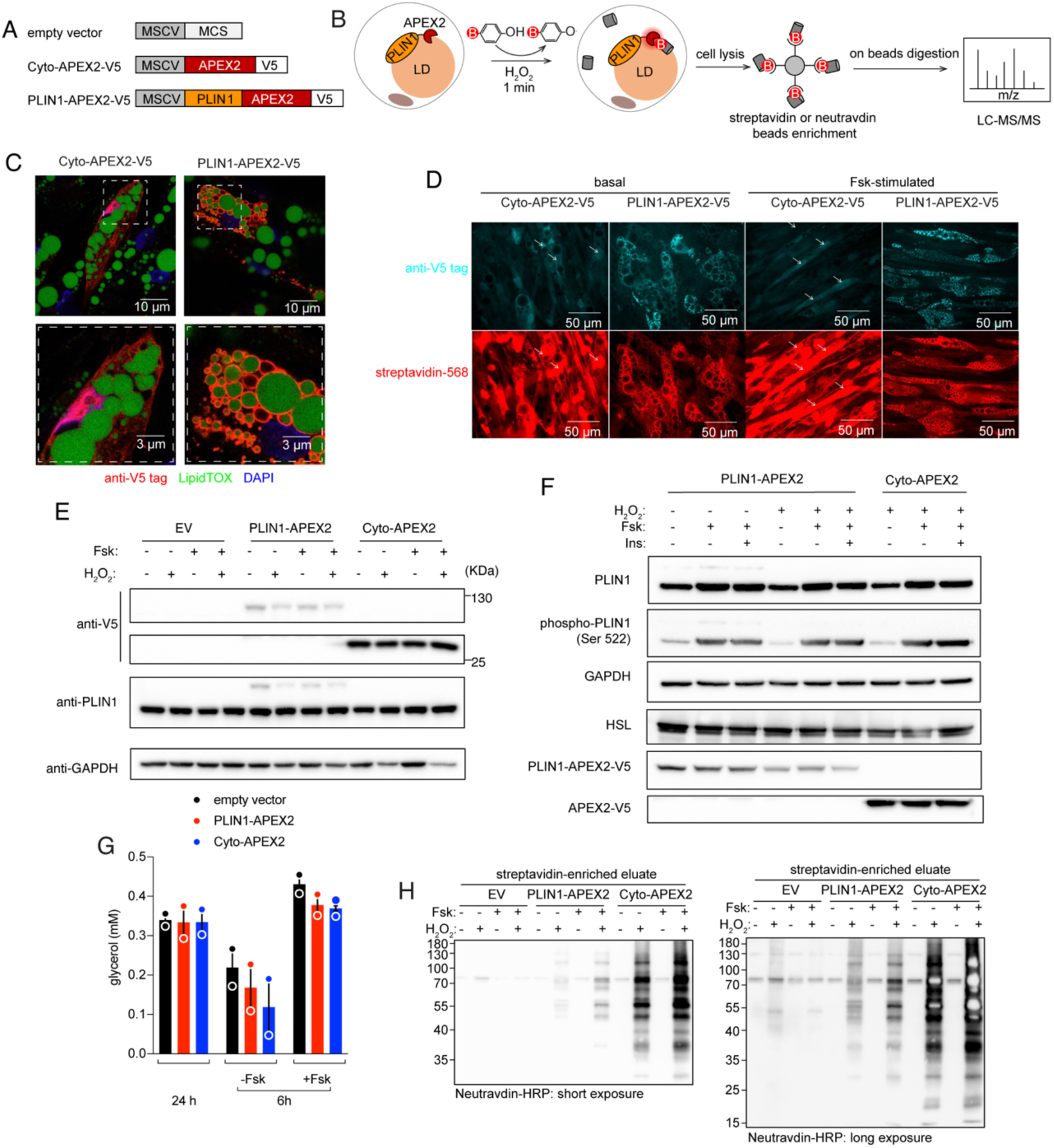
PLIN1-APEX2 fusion construct targets to LDs and maintains enzymatic activity without perturbing lipolysis in human adipocytes. (A) Lentiviral vectors used in this study. (B) Workflow of PLIN1-APEX2-mediated proximity labeling. (C) Images of human primary adipocytes transduced with Cyto-APEX2-V5 and PLIN1-APEX2-V5, stained for V5 epitope tag (red), LipidTOX (green) and DAPI (blue). Magnified regions below highlight the cellular regions with lipid droplets. (D) Cells treated with biotin-phenol/hydrogen peroxide stained for V5 epitope tag (cyan) and with streptavidin-HRP (red) to detect the biotinylated proteins. (E,F) Western blots of whole-cell lysates from cells treated as described above each label, probed as indicated to the left of each band. (G) Glycerol concentration in cell medium (error bars = range, n = 2, biological replicates). (H) Western blot of proteins enriched by streptavidin beads, visualized using neutravidin-HRP for short (left) and long (right) exposures.

Since overexpression of proteins can lead to perturbation of their physiological functions, we evaluated the level of expressed constructs. Western blotting of whole-cell lysates using anti-V5 antibody revealed bands at the expected molecular weights of PLIN1-APEX2-V5 (~90 kDa) and Cyto-APEX2-V5 (~27 kDa) in cells expressing the corresponding constructs, with significantly more abundant Cyto-APEX2-V5 expression (Fig. 1E). Probing with anti-PLIN1 antibody revealed a band at the expected molecular weight of endogenous PLIN1 (~ 60 kDa) in all samples, and an additional, much fainter band corresponding to PLIN1-APEX2-V5 in cells transfected with the corresponding construct. Thus, PLIN1-APEX2-V5 was expressed at much lower levels than endogenous PLIN1. To verify that expressed PLIN1-APEX2-V5 did not functionally interfere with endogenous PLIN1, we measured basal and Fsk-stimulated PLIN1 phosphorylation and lipolysis. We observed similarly enhanced phosphorylation of PLIN1 in response to Fsk in cells expressing both constructs (Fig. 1F). Moreover, PLIN1 phosphorylation was similar in cells expressing PLIN1-APEX2-V5 incubated without or with H_2_O_2_, indicating that the conditions employed during the biotinylation reaction did not interfere with signal transduction (Fig. 1F). To monitor lipolysis, we measured basal glycerol accumulation in the media over 48h, and glycerol release over 6 h in the presence or absence of Fsk. No difference in glycerol accumulation was observed between cells expressing empty vector, PLIN1-APEX2-V5 or Cyto-APEX2-V5 (Fig. 1G).

We then probed the western blots with HRP-streptavidin to obtain an overview of all proteins biotinylated by each construct. In lysates of cells transduced with empty vector, only few faint bands, possibly corresponding to endogenous biotin-dependent carboxylases were detected. In contrast, a large number of biotinylated proteins were seen in lysates from Cyto-APEX2-V5 and PLIN1-APEX2-V5 transduced cells (Fig. 1H), with significantly more in the former condition.

### Identification of biotinylated proteins by LC-MS/MS

We obtained biotinylated proteins from cells transduced with PLIN1-APEX2-V5, Cyto-APEX2-V5, or cells transduced with PLIN1-APEX2-V5 but not exposed to H_2_O_2_ therefore preventing biotinylation (Negative Control). Transduced cells were treated without or with Fsk for 6h, and one set of cells was treated with insulin for 1h prior to addition of Fsk (Fig. 2A). Biotinylated proteins were recovered from three independent sets of transduced cells and identified by LC-MS/MS following on-beads digestion. The number of proteins identified (LFQ > 0 in > 5 out of 9 replicates) was on average 205, 402 and 481 in the Negative Control, PLIN1-APEX-V5 and Cyto-APEX2-V5 groups, respectively (Fig. 2B, Supplementary Table S1). Principal component analysis (PCA) revealed three distinct clusters, but different lipolytic conditions within each cluster (i.e., presence of Fsk or Fsk+insulin) did not significantly contribute to the variance (Fig. 2C).

**Figure 2.**
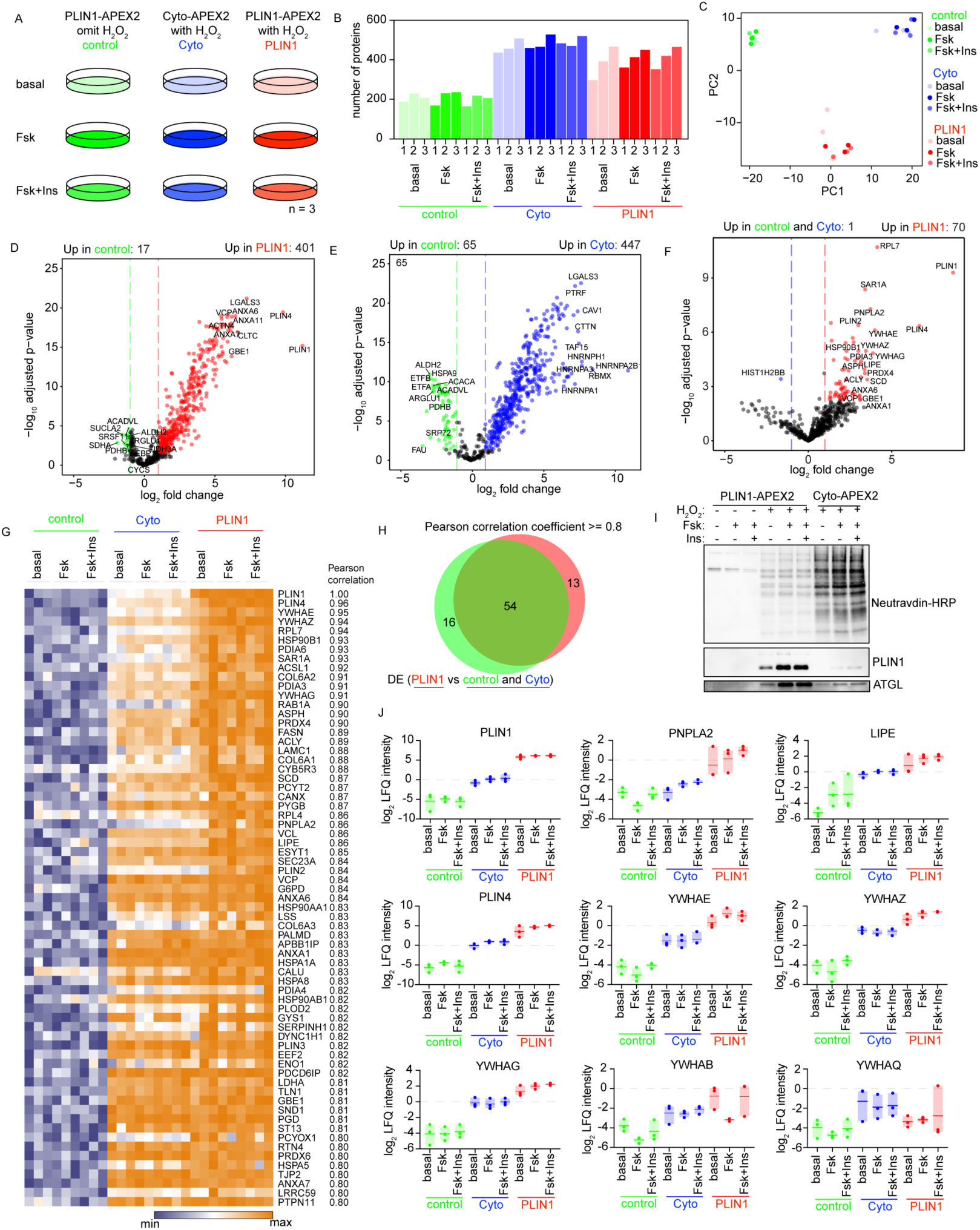
Identification of LD-associated proteins. (A) Design of labeling and proteomic experiments. Results are derived from three independent experiments. (B) Bar graph showing the total number of proteins identified by LC-MS/MS in each sample. (C) Scatter plot of the first two principal components of LFQ intensities of proteins identified by LC-MS/MS across all the samples. (D,E) Volcano plots of biotinylated proteins enriched across all samples (Basal, Fsk and Fsk+Ins) by PLIN1-APEX2 (D) or Cyto-APEX2 (E) compared to negative control samples. (F) Volcano plot of biotinylated proteins enriched across all samples (Basal, Fsk and Fsk+Ins) by PLIN1-APEX2 compared to combined negative control and Cyto-APEX2. Colored symbols in D, E and F correspond to Benjamini-Hochberg-adjusted p-value < 0.05 and log2 fold change ≥ 1.0. (G) Heatmap of proteins enriched by PLIN1-APEX2 based on Pearson correlation coefficient > 0.80 with PLIN1 values across all samples. (H) Venn diagram showing the overlap between LD proteomes identified by differential enrichment analysis (F) and Pearson correlation analysis (G). (I) Western blot analysis using neutravidin-HRP and antibodies against PLIN1and ATGL for proteins enriched by neutravidin beads. (J) Box plots of LFQ intensities of indicated proteins in n=3 independent replicate samples.

Differential enrichment analysis using LFQ intensity values (Benjamini-Hochberg-adjusted p-value < 0.05 and log2 fold change ≥ 1.0) revealed 401 proteins to be significantly enriched by PLIN1-APEX2-V5 compared to Negative Control (Fig. 2D, Supplementary Table S2, Tab 1), with the highest differentially expressed proteins corresponding to PLIN1 and PLIN4. 447 proteins were significantly enriched by Cyto-APEX2-V5 relative to Negative Control (Fig. 2E, Supplementary Table S2, Tab 2). 70 proteins were significantly enriched by PLIN1-APEX2-V5 relative to the combined Negative Control and Cyto-APEX2-V5 samples (Fig. 2F, Supplementary Table S2, Tab 3). Among these 70 proteins, PLIN1 PLIN2, PLIN4, RAB1A, RAB7A, CYB5R3, PNPLA2, LIPE, VCP, and LSS have been previously reported to be associated with lipid droplets. We searched for proteins specifically enriched by PLIN1-APEX2-V5 in response to Fsk or Fsk+Insulin. Unexpectedly, neither Fsk or Fsk+Ins treatment resulted in significant changes in the basal PLIN1interactome (Fig. S1, S2, Supplementary Table S2, Tabs 4-8), suggesting that the regulation of lipid mobilization might be more dependent on activities of key proteins rather than their abundance on the LD surface. To identify all potential PLIN1-interacting proteins, we analyzed enrichment across all lipolytic conditions. Proteins were ranked by the correlation of their imputed LFQ intensity values with those of PLIN1 (Pearson correlation coefficient >= 0.8) (Fig. 2G, Supplementary Table S3). Proteins correlated with PLIN1 included known LD binding proteins and multiple additional ER proteins, and largely overlapped with those found by differential enrichment analysis (Fig. 2H). To verify the results obtained by LC-MS/MS, we conducted western blotting of recovered biotinylated proteins, and observed significant enrichment of PLIN1 and ATGL by PLIN1-APEX2-V5 (Fig. 2I).

We were intrigued by the enrichment of four members of the 14-3-3 family of proteins (YWHAB, YWHAE, YWHAG and YWHAZ) by PLIN1-APEX2-V5, as these are adapter/scaffold proteins that bind to phosphorylated targets, regulating their activity, localization, and/or stability. Moreover, YWHAB interaction with PLIN1 was decreased by Fsk (Supplementary Table S2, Tab 4), and this decrease was reversed by insulin (Supplementary Table S2, Tab 6), although these interactions did not reach statistical significance upon adjustment for multiple comparisons. We compared the LFQ intensity values of all detected 14-3-3 family proteins (YWHAE, YWHAZ, YWHAG, YWHAB, and YWHAQ) to those of the known lipid droplet proteins PLIN1, PNPLA2, LIPE and PLIN4 (Fig. 2J). Three of these proteins (YWHAE, YWHAZ, and YWHAG) are enriched by PLIN1-APEX2 under all three lipolytic conditions, while YWHAB seemed more variable, being enriched only under basal and Fsk+Insulin conditions.

To shed light on the potential biological significance of the PLIN1 interactome in human adipocytes, we contextualized our results by comparing them with those obtained by Bersuker et al (19), who used APEX2 to identify LD proteins in two cancer cell lines (U2OS and Huh7). Using a combination of PLIN2-APEX2 and ATGL-APEX2, Bersuker et al. identified 1302 and 699 total biotinylated proteins in U2OS and Huh7 cells, compared to our 599 proteins identified in human adipocytes by PLIN1-APEX2. 174 proteins were identified in all three cell types (Fig. 3A, Supplementary Table S4). Of these, Bersuker et al. identified 57 proteins as associated with LDs with high confidence, close in number to the 54 proteins identified in our study (Fig. 2G). Of the high-confidence LD proteins, seven proteins (PLIN2, PLIN3. PNPLA2, RAB1A, CYB5R3, LSS and VCP) were identified in all three cell types (Fig. 3B, Supplementary Table S4), suggesting they represent the core machinery necessary for LD assembly.

**Figure 3.**
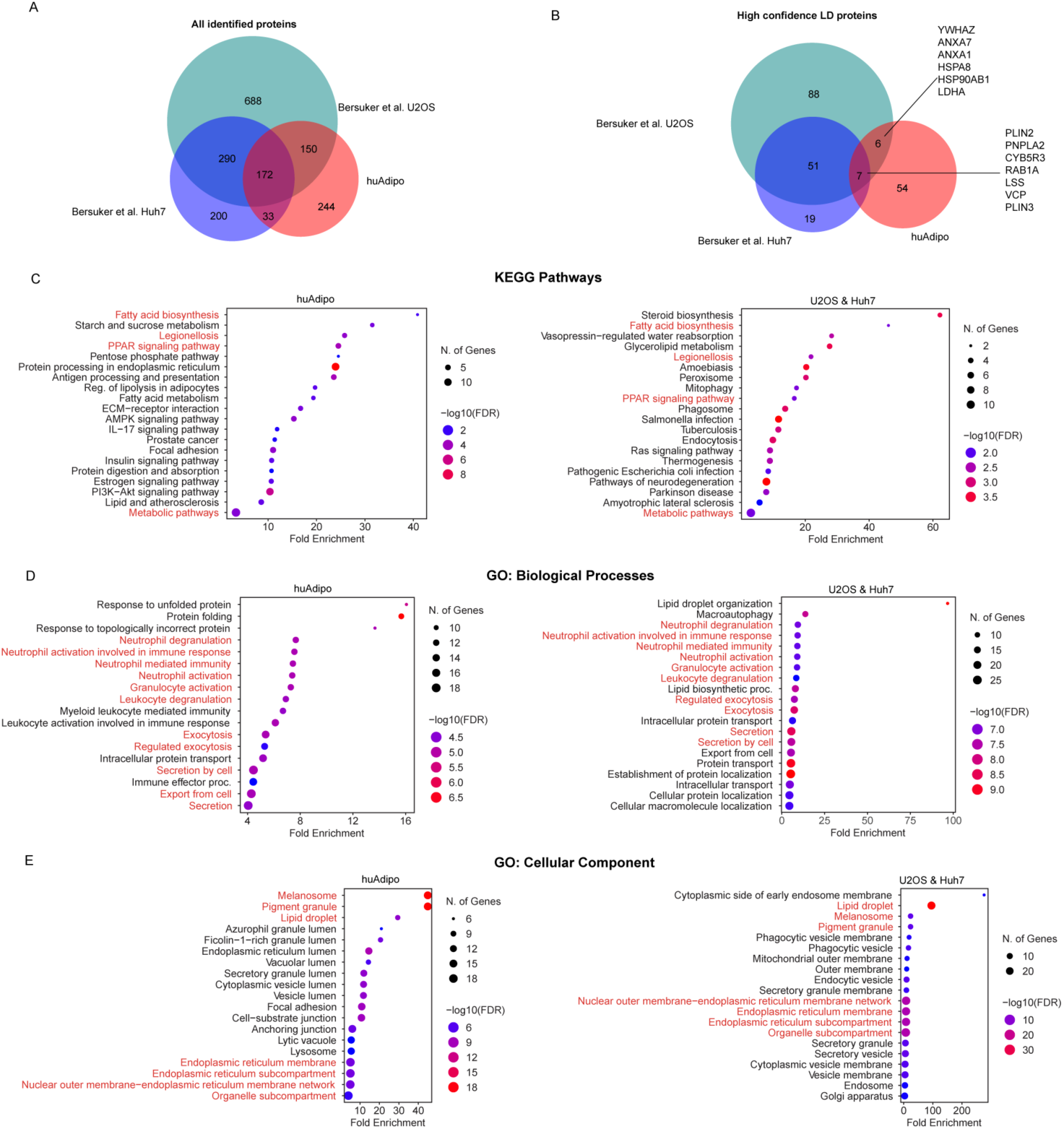
Comparison of LD proteins identified in human primary adipocytes and other cell lines. (A,B) Venn diagrams of all proteins (A) and high-confidence LD proteins (B) identified in U2OS and Huh7 by Bersuker et al. (19), and human primary adipocytes. (C,D,E) KEGG pathway analysis (C), GO biological process terms (D), and GO Cellular Component terms (E), of high confidence LD proteins from human primary adipocytes (left) and from U2OS/Huh7 cells (right). Pathways in common are highlighted in red.

To distinguish aspects of LD function that might be cell type-specific, we conducted pathway analyses on the high confidence LD proteins from human adipocytes (Left panels, Fig. 3) or U2OS and Huh7 (Right panels, Fig. 3). Pathway analysis (KEGG) showed common enrichment in pathways of fatty acid biosynthesis and PPARψ signaling, but pathways including AMPK signaling, IL-17 signaling and PI3K-Akt signaling were enriched only in adipocyte LDs (Fig. 3C, Supplementary Table S5, Tabs 1,2). Gene ontology defined biological processes in both cases were enriched for immune cell functions, involving membrane trafficking proteins and chaperones (Fig. 3D, Supplementary Table S5, Tabs 3,4), and consistent with a proposed role for lipid droplets in early innate immune responsiveness (21, 22). Cellular components identified as enriched by high-confidence LD proteins also included the endoplasmic reticulum, lysosomes, and mitochondria (Fig. 3E); 11 proteins detected in human adipocyte LDs are associated with mitochondria, 36 with the endoplasmic reticulum, 28 with the nucleus and the remainder are cytoplasmic or of indetermined subcellular localization. These results are consistent with the existence of extensive contact sites between LDs and other organelles in all other cell types (23). In aggregate, the contrasting characteristics of LD proteins in human adipocytes versus cancer cell lines suggest that signaling pathways associated with systemic metabolic control play a more substantial role in the regulation of adipocyte LDs.

### Functional role of 14-3-3 proteins

To verify the presence and further explore the function of the 14-3-3 proteins enriched on adipocyte lipid droplets, we first analyzed the specificity of commercially available antibodies for this family of proteins. We generated HA-tagged versions of YWHAB, YWHAG, YWHAZ and YWHAE, expressed them in 293T cells, isolated the proteins by immunoprecipitation using anti-HA antibodies, and conducted western blotting. Most commercially available antibodies lacked specificity, except for those raised against YWHAG and one to YWHAB (Fig. S3). Western blotting of proteins biotinylated by PLIN1-APEX2-V5 using an antibody that detects YWHAB, G, E and Z (Abcam) revealed enrichment relative to Cyto-APEX2-V5 under all treatment conditions (Fig. 4A). Immunofluorescence analysis of adipocytes using the same antibody revealed a punctate pattern surrounding the lipid droplets, consistent with close interactions of 14-3-3 proteins with PLIN1 on the LD surface (Fig. 4B).

**Figure 4.**
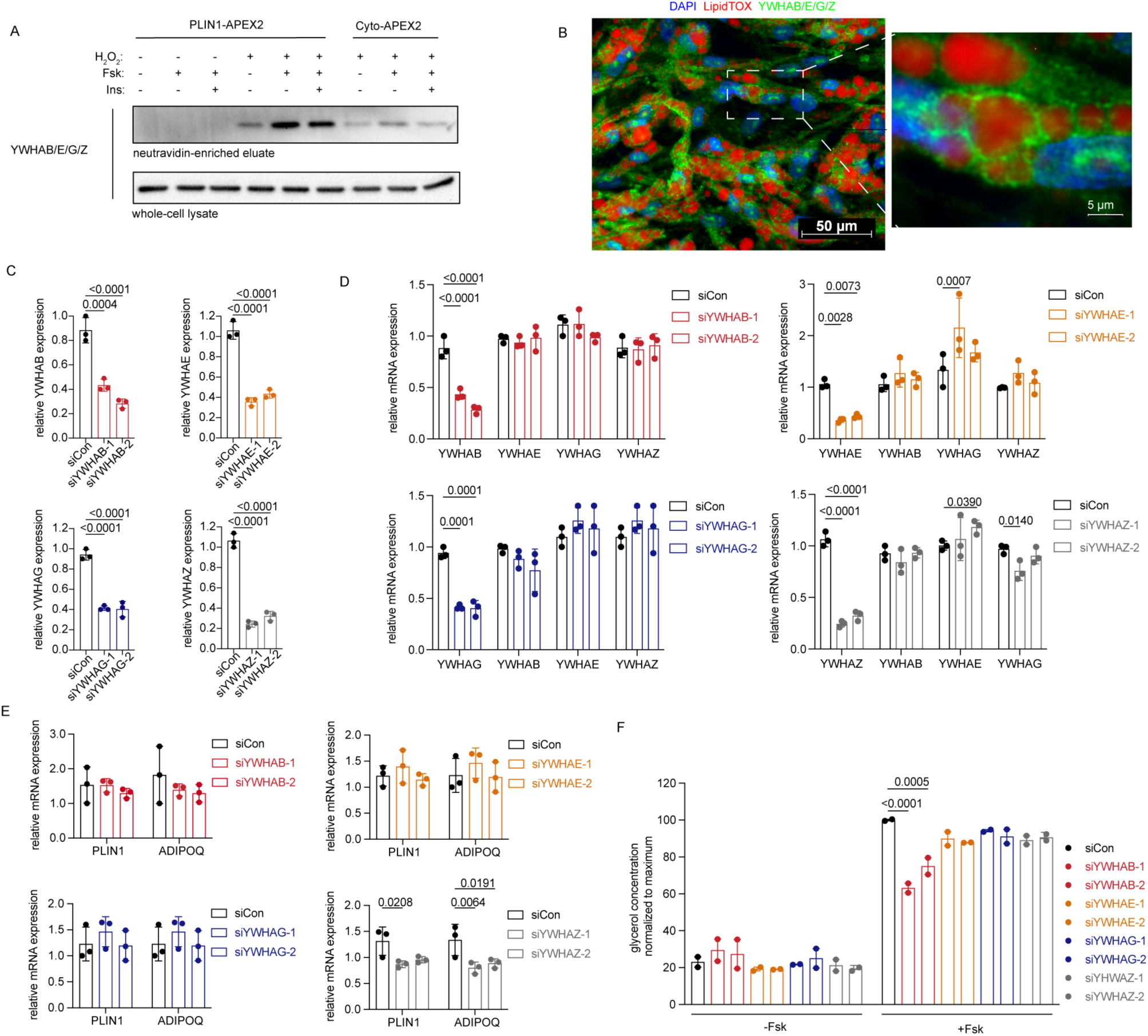
Analysis of 14-3-3 proteins. (A) Western blot probed with antibody recognizing YWHAB/Z/G/E of proteins biotinylated by PLIN1-APEX2 enriched by neutravidin beads compared to whole-cell lysate. (B) Immunofluorescence analysis of primary adipocytes using antibody recognizing YWHAB/Z/G/E (green), DAPI (blue) and LipidTOX (red). (C) RT-PCR of RNA obtained 48h after transfection of adipocytes with two siRNAs targeting *YWHAB*, *YWHAE*, *YWHAZ* and *YWHAG*, as indicated in the Y axis. (D) RT-PCR of RNA obtained 48h after transfection of adipocytes transfected with two siRNAs targeting *YWHAB*, *YWHAE*, *YWHAZ* or *YWHAG*, as indicated in the legend, probed for the transcript indicated in the X-axis. (E) RT-PCR analysis of RNA obtained 48h after transfection of adipocytes with siRNAs targeting *YWHAB*, *YWHAE*, *YWHAZ* and *YWHAG*, probed for expression of *PLIN1* and *ADIPOQ.* For C,D and E, values are expressed as the fold relative to siRNA control values. Bars represent means and SD of n = 3 biological replicates. Statistical differences were assessed using two-way ANOVA, and exact P values are shown. (F) Glycerol accumulation over 6h from cells treated with vehicle or Fsk, 48h after transfection with siRNAs targeting *YWHAB*, *YWHAE*, *YWHAZ* and *YWHAG.* Bars represent the mean and SD of n = 2 independent experiments, each conducted with 3-fold technical replication. Statistical differences were assessed using two-way ANOVA, and exact P values are shown.

We then used siRNAs to knock down the expression of each of the LD enriched 14-3-3 proteins in differentiated adipocytes. We achieved 60~70% knockdown efficiency for each isoform (Fig. 4C), and found that knockdown of the targeted isoforms did not alter the expression level of the non-targeted isoforms (Fig. 4D), consistent with absence of compensatory effects. To determine whether knockdown of these genes would influence adipocyte differentiation, we measured expression levels of PLIN1 and ADIPOQ by RT-PCR (Fig.3E). While knockdown of YWHAB, YWHAE and YWHAG had no significant effect, YWHAZ knockdown significantly decreased differentiation (Fig. 4E). To assess specific LD functions, we measured basal and stimulated lipolysis (Fig 4F). Basal lipolysis was not affected by the knockdown of any of the isoforms, but Fsk-stimulated lipolysis was decreased specifically in response to knockdown of YWHAB (Fig. 4F).

To further explore the functional role of YWHAB we employed CRISPR-Cas9 to achieve full deletion of the protein. To define the optimal genomic locus for YWHAB gene disruption, we designed 4 sgRNAs: sg-YWHAB-1 targeted exon 5 and sg-YWHAB-2, sg-YWHAB-3, and sg-YWHAB-4 targeted exon 3, both exons within the open reading frame of the YWHAB gene (Fig. 5A). We delivered the RNP complexes containing Cas9 and sgRNA to progenitor cells according to a previously developed method(24). Small insertions and deletions were generated by the 4 different sgRNAs in 85-99% of cells (Fig. 5B), and the deletions were largely maintained upon differentiation of progenitors into adipocytes (83-87% editing efficiency) (Fig. 5C). To determine whether the deletions would result in protein deficiency, we performed western blotting with verified antibodies to YWHAB and, for comparison, YWHAG. While YWHAB was undetectable under all lipolytic conditions tested, no difference in YWHAG expression was observed (Fig. 5D). The abundance of PLIN1 was not affected by YWHAB deficiency (Fig. 5D), consistent with phase imaging of lipid droplets indicating that YWHAB knockout did not cause the impairment of cell differentiation (Fig. 5E). Notably, cAMP-stimulated signal transduction to PLIN1 was not affected by YWHAB deficiency, as the effects of Fsk to stimulate PLIN1 phosphorylation were not impaired (Fig. 5D). The efficient CRISPR-Cas9 knockdown of YWHAB allowed us to verify its proximity to PLIN1 using proximity ligation (Fig. 5 F and G). A clear signal reflecting the proximity of anti-PLIN1 and anti-YWHAB antibodies was detected surrounding the lipid droplets (Fig. 5F, top panels), and this signal was abrogated in cells lacking YWHAB (Fig. 5F, bottom panels), as quantified by image analysis (Fig. 5G)

**Figure 5.**
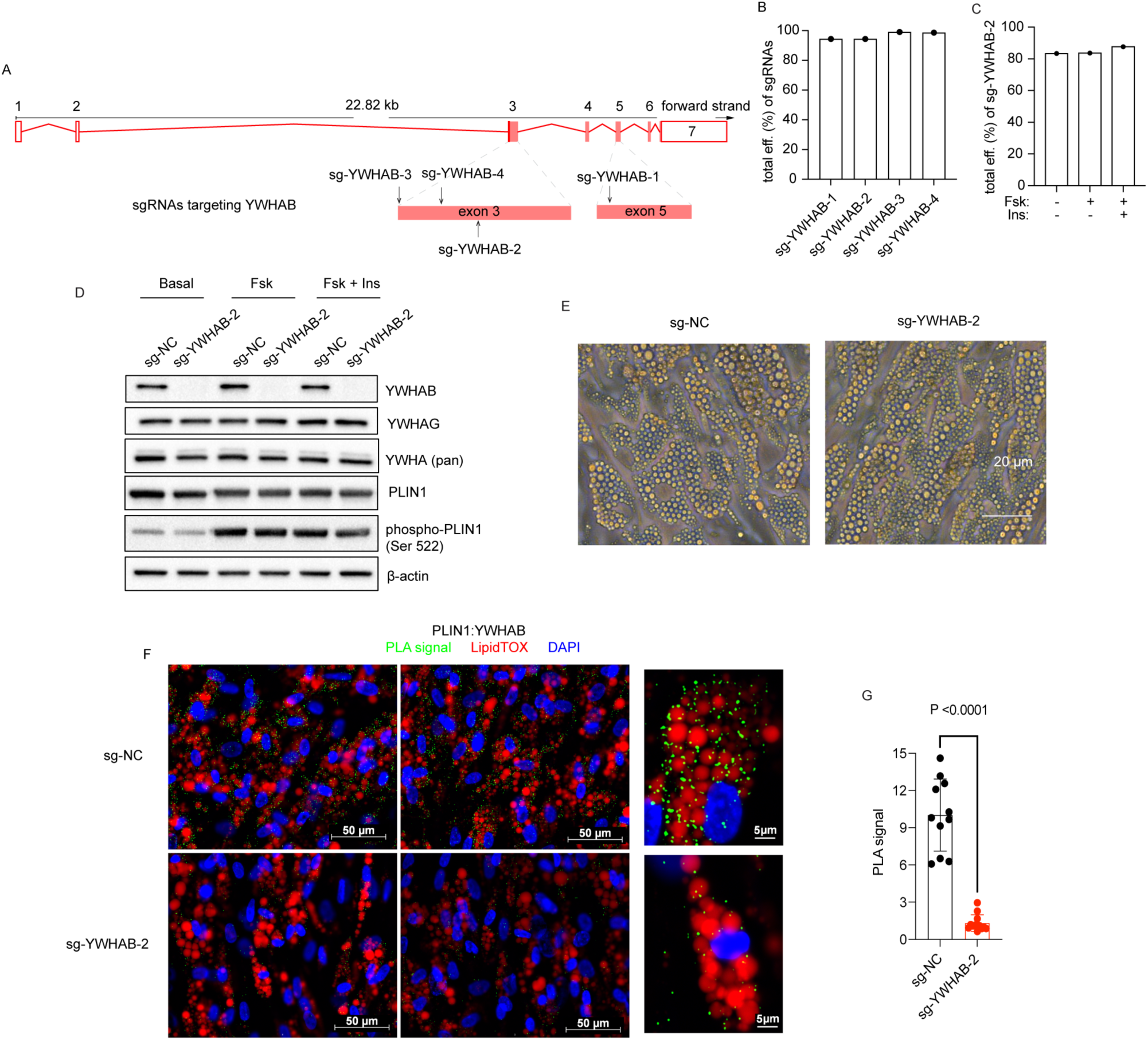
CRISPR-Cas9 mediated knockout of YWHAB. (A) Localization of sg-YWHAB-1, sg-YWHAB-2, sg-YWHAB-3, and sg-YWHAB-4 in exons 3 and 5 of the human YWHAB coding region. (B,C) Editing efficiency of sgRNAs in progenitor cells (B) and differentiated adipocytes (C), determined by TIDE analysis. (D) Western blots for proteins indicated from adipocytes differentiated from progenitor cells electroporated with non-targeting control (sg-NC) or sg-YWHAB-2 under different lipolytic conditions. (E) Representative phase images of human adipocytes without or with YWHAB knockout on day 8 of differentiation. (F) Representative images of adipocytes differentiated from cells electroporated with non-targeting control (sg-NC, Top panels) or sg-YWHAB-2 (Bottom panels), visualizing the proximity ligation product (PLA, green) of antibodies to PLIN1 and YWHAB. Cells are counterstained with DAPI (blue) and LipidTOX (red). (G) Mean and SD of total intensities of PLA signal in n=11 independent fields of cells stained as in (F). Statistical difference was assessed using two-tail student t-test, and exact P value is shown.

We then examined the functional effects of YWHAB knockout on lipolysis in cells derived from two independent donors (Fig. 6, A-F). Fsk induced a dose-dependent increase in glycerol accumulation, resulting from triglyceride lipolysis, which was observed after 6 h (Fig. 6, A-C) and persisted for 24h (Fig. 6 D-F) after stimulation. Insulin mitigated the lipolytic effects of Fsk in both cell lines, and this effect was most prominent at the 6h time points (Fig. 6, A-C). Fsk-stimulated glycerol accumulation was impaired by YWHAB knockout in both cell lines at 6h (Fig. 6, A-C), but was less evident after 24h (Fig. 6, D-F). Notably, the anti-lipolytic effect of insulin was enhanced in cells lacking YWHAB at both time points (Fig. 6, C,F). To determine whether the effects of YWHAB knockout might be attributable to changes in cell viability, we measured ATP levels (Fig. 6, G-I). We find that Fsk treatment decreases ATP levels at all concentrations tested, but this decrease was not enhanced by YWHAB knock out. (Fig. 6I). These results indicate that YWHAB is required for optimal stimulation of lipolysis, and that the anti-lipolytic effect of insulin is suppressed by YWHAB.

**Figure 6.**
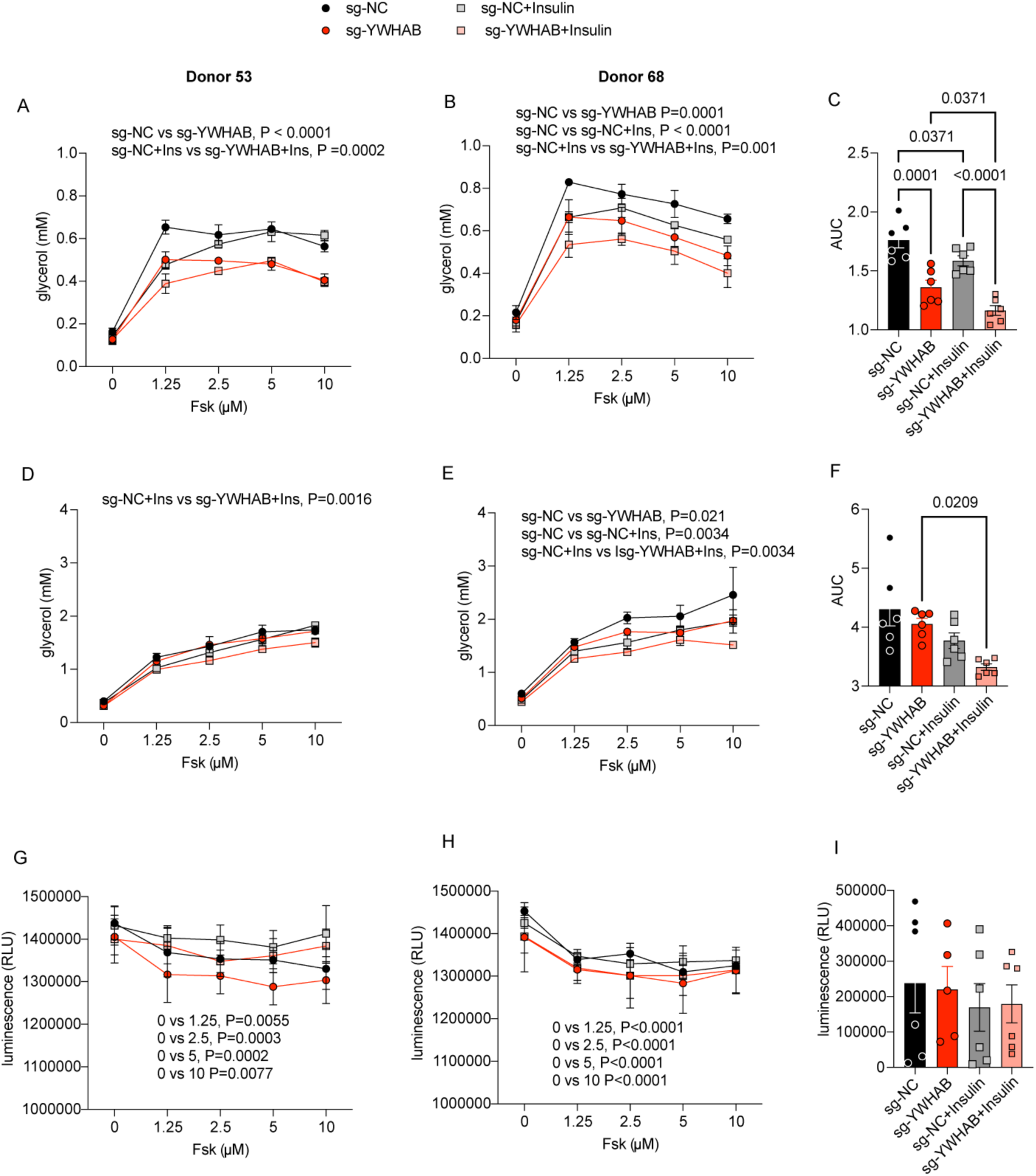
Functional effects of YWHAB deletion. Adipocytes differentiated from cells electroporated with non-targeting control (sg-NC) or sg-YWHAB-2, were stimulated with different concentrations of Fsk without or with the addition of insulin. Glycerol accumulation was measured after 6 (A-C) and 24 (D-F) hours of stimulation, and CellTiter-Glo 2.0 assay was performed after 24 h of stimulation. Shown are means and SD of values derived from n=3 independent cultures of cells derived from two donors (donor 53, A,D,G; donor 68, B,E,H), and area under the curve of the combined results from both donors (C,F,I). Statistical significance was determined using two-way (A,B,D,E,G,H) or one-way (C,F,I) ANOVA, and exact P values are shown.

## DISCUSSION

Lipid droplets are found in all cells, where they play a key energetic role by sequestering and releasing lipids for intracellular use. Adipocyte lipid droplets have an additional, key function of meeting the energetic needs of the entire organism. It is therefore of great interest to understand the similarities and differences between non-adipocyte and adipocyte LD function and how specific adipocyte LD mechanisms might operate and impact metabolic health. Prior research has shown that many of the molecular mechanisms controlling LD dynamics are mediated by specialized proteins that target the LD surface, such as the perilipins. In this study we define proteins that are in close proximity to PLIN1 in human adipocytes, and by comparison to those discovered in other cells by similar approaches, conclude that adipocyte LDs are enriched in proteins associated with hormonal signaling. In particular, multiple isoforms of the 14-3-3 family of proteins are enriched in the vicinity of PLIN1, and we find that one of them, YWHAZ, is necessary for adipocyte differentiation, and another, YWHAB, enhances lipolysis and mitigates the anti-lipolytic actions of insulin.

The comparison of proteins interacting with PLIN1 in adipocytes with those interacting with PLIN2 and ATGL in two cancer cell lines provides insight into core components of LDs. Of all identified proteins, only 7 (PLIN2, PLIN3, PNPLA2, CYB5R3, RAB1A, LSS, VCP) are found in all three cell types. PLIN1 and PLIN2 are known perilipins, and PNPLA2/ATGL is the major mediator of hormonally stimulated lipolysis. CYB5R3, RAB1A, LSS and VCP have all been localized to the endoplasmic reticulum, highlighting the close relationship between LD function and biogenesis – indeed, PLIN1 has also been reported to localize to the ER in some studies (25), and LDs and ER maintain close contacts (23). Beyond these 7 core common LD proteins, many others are associated with lipid synthetic pathways, as expected given the core function of LDs as reservoirs of triglycerides and other lipids. Interestingly, LD proteins also include chaperones and trafficking proteins that operate in exocytosis, which are enriched in immune cell pathways. These associations suggest that LDs may have an important role in immune cell function, but also suggests that exocytosis of lipids via vesicular trafficking may be a central, less recognized role of LDs (26). In addition to proteins common to multiple cell types, our study identifies proteins more specifically associated with adipocyte LDs, mapping to the PI-3 kinase, AMP kinase, insulin, estrogen and IL-17 signal transduction pathways. These results indicate that adipocyte LDs are poised to respond to hormonal signaling to rapidly meet systemic energy needs.

While proximity labeling is more specific than traditional biochemical methods and capable of capturing LD proteome in intact cells, there are still some limitations in our study. Not all previously reported LD proteins, notably ABHD5 (27), were identified by PLIN1-APEX2 labeling, nor by PLIN2-APEX2 by Bersuker et al. (19). Moreover, we could not determine statistically significant differences in proteins labeled by PLIN1 under different lipolytic conditions, despite clear effects on glycerol release. The absence of major differences may be attributable to our use of label-free proteomics, which may be less quantitative compared to other techniques such as stable isotope labeling by amino acids in cell culture (SILAC) (18) or tandem mass tags (TMT) (28). Alternatively, lipolysis may be more dependent on activities of LD proteins, and less to architectural rearrangements of the LD proteome. Despite these limitations, our analysis revealed significant enrichment by PLIN1-APEX2 of proteins implicated in signal transduction, such as YWHA (14-3-3) proteins.

The mammalian YWHA/14-3-3 protein family is comprised of seven isoforms encoded by distinct genes. They preferentially exist as homodimers but can also heterodimerize; each monomer binds to serine/threonine phosphopeptide motifs with an ordered affinity ranking (29), protecting from de-phosphorylation. By binding two phosphopeptides, 14-3-3 dimers can change the conformation of a phosphoprotein or mediate its interaction with other phosphoproteins, thereby modulating localization, enzymatic activity and multiple cellular functions (30). A major role of 14-3-3 proteins in the maintenance of cellular architecture has recently been proposed (31).

In the context of adipocyte function, 14-3-3 proteins have been reported to interact with ACACA, ACLY, and FASN (32), suggesting a key role in the regulation of fatty acid biosynthesis. YWHAZ has been implicated in adipocyte differentiation, potentially through effects on RNA splicing (33–35), and consistent with our finding that knockdown of YHWAZ, but not other isoforms, impairs human adipocyte differentiation. Interestingly, YWHAB has been implicated in insulin signaling, through interactions with the insulin receptor, IRS1 and also phosphodiesterase 3B. (12, 36, 37). In our study, complete ablation of YWHAB in human adipocytes did not impair insulin-mediated suppression of lipolysis; indeed, maximal suppression by insulin was seen in cells lacking YWHAB. Together these studies suggest that interaction of YWHAB with elements of the insulin signaling pathway may overall downregulate insulin signaling.

The enrichment of four isoforms, YWHAE, YWHAZ, YWHAG and YWHAB, in the vicinity of PLIN1, as we infer from their biotinylation by PLIN1-APEX2-V5, is consistent with a major role of phosphorylation/dephosphorylation reactions in regulation of LD function. Current models of lipolysis activation postulate that cAMP-dependent protein kinase A is indispensable for activation of PLPLA2/ATGL by ABHD5, which is sequestered by PLIN1 under basal conditions and released following PLIN1 phosphorylation. Additionally, ATGL and HSL are also phosphorylated. The timing and precise localization of these events on the LD surface may be coordinated by the actions of the YWHA proteins in the vicinity of PLIN1, are consistent with identification of YWHAQ and YWHAZ by proximity labeling with PLIN2 (19) and of YWHAZ/G/E/Q though biochemical methods (38, 39), Our results advance our understanding of the functional role of YWHAB by demonstrating its requirement for maximal cAMP-stimulated lipolysis and its ability to mitigate the anti-lipolytic effects of insulin. Further studies will be required to identify the specific molecular interactions mediated by YWHAB, and of how these interactions may advance the development of therapeutic interventions for metabolic diseases.

## MATERIALS AND METHODS

### Primary progenitor cells

Progenitor cells were obtained from human abdominal subcutaneous adipose tissue explants and expanded according to previously published methods (18). Briefly, adipose tissue explants from patients undergoing panniculectomy procedures at University of Massachusetts Medical Center with the approval of University of Massachusetts Institutional Review Board (#14734_13) were embedded in MatriGel (Corning, cat# 356231), (200 1 mm^3^ explants per 10 cm dish) and cultured for 14 days. Progenitor cells sprouting from the explants were obtained using Dispase followed by Trypsin-EDTA and Collagenase I treatment and expanded in 15-cm cell culture dishes. To induce adipogenic differentiation, progenitor cells at 100% confluency were maintained in DMEM + 10% FBS supplemented with 0.5 mM 3-isobutyl1-methylxanthine, 1 µM dexamethasone and 1 µg/ml insulin (MDI) for 72 hrs. Subsequently, half of the medium was replaced with DMEM + 10% FBS every other day for 3 days. On day 8 of differentiation, cells were treated with forskolin (10 µM) without or with insulin (5 µg/ml) for 6 hrs. For immunofluorescence staining experiments, cells were grown on glass coverslips in 24-well plate. For qPCR experiments, cells were grown in 24-well plates. For small-scale enrichment of biotinylated proteins, cells were grown on 6-well plates. For labeling experiments followed by mass spectrometry analysis, cells were grown on 10-cm dishes. For western blot analysis, cells were grown in 12-well plates.

### PLIN1-APEX2-V5 and Cyto-APEX2-V5 constructs

PLIN1 and APEX2-V5 fragments were amplified by PCR from PLIN1 cDNA ORF clone in pcDNA3.1^+^/C-(K)-DYK vector (GenScript: OHu22113) and Twinkle-APEX2-V5 vector (Addgene, plasmid #129705) respectively. PLIN1-APEX2-V5 construct was generated by assembling PLIN1 and APEX2-V5 fragments into lentiviral vector pCDH-MSCV-MCS-EF1-GFP by HiFi DNA assembly. GGGGS was added as a fusion protein linker between PLIN1 and APEX2-V5. Cyto-APEX2-V5 construct was generated by assembling APEX2-V5 fragment into the same vector by HiFi DNA assembly. Both constructs were validated by sanger sequencing. The primers used for PCR amplification and sequencing were listed below.

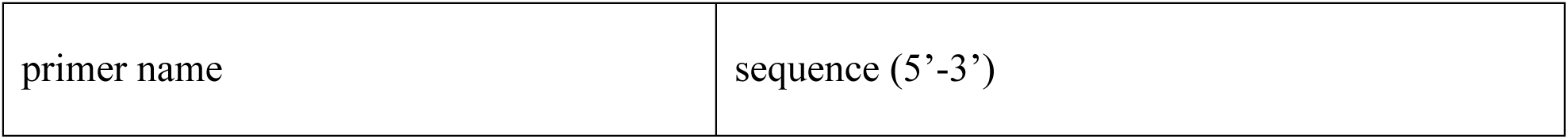
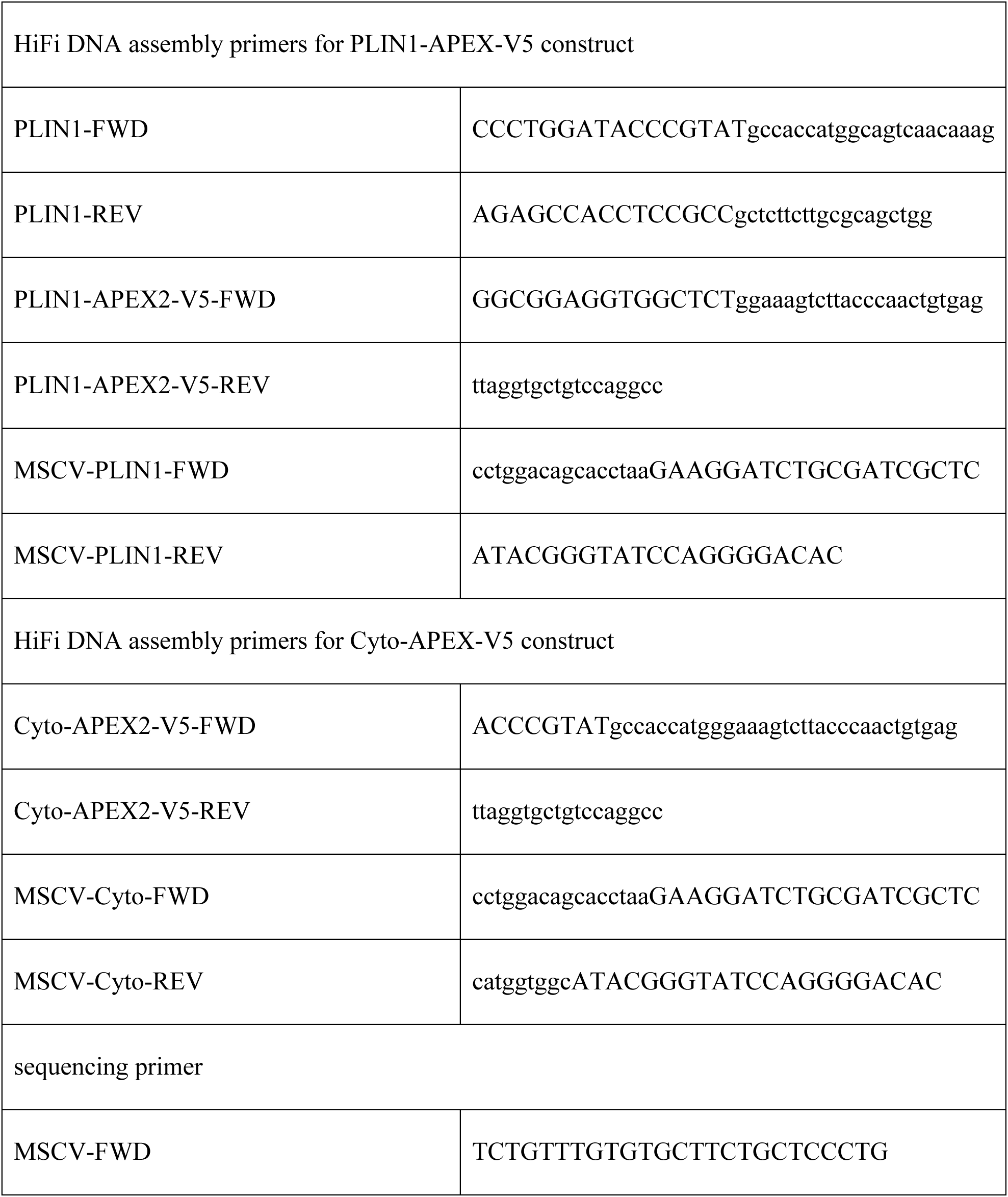

### Lentivirus production

Lenti-X 293T cells in 15-cm dishes were transfected at ~70% confluency with the lentiviral vector pCDH-MSCV-MCS-EF1-GFP containing gene of interest (18 µg), packaging plasmid psPAX2 (13.5 µg) and envelop plasmid pMD2.G (4.5 µg) using calcium phosphate transfection. 48 and 72 hours after transfection, the medium containing lentiviruses was centrifuged at 500 x g for 10 min at 4°C and filtered through a 0.45-mm filter to remove cells and debris. The filtered lentiviral supernatants were concentrated using Lenti-X concentrator according to the manufacturer’s instructions. The concentrated lentiviruses were titrated by abm qPCR lentivirus titer kit. Afterwards, they were used for human adipocytes transduction immediately or stored at −80°C in single-use aliquots.

### Lentiviral transduction of human adipocytes

Cells were grown in 10-cm dishes as described above, and on day 5 of differentiation transduced with the volume of empty vector, PLIN1-APEX2-V5 or Cyto-APEX2-V5 lentiviruses at a MOI of 50 in the presence of 8 µg/ml polybrene. 24 hours after transduction, the lentiviral supernatant was replaced with fresh complete DMEM. The cells were maintained in DMEM for two more days until APEX labeling on day 8 of differentiation.

### In vivo APEX2 labeling in human adipocytes

On day 8 of differentiation, cells were treated with 10 µM forskolin (Fsk) (Sigma, cat# F3917) without or with insulin (5 µg/ml) for 6 hours. To initiate APEX2 labeling, cells were incubated with 3 ml of 500 µM biotin-phenol in complete DMEM (containing the corresponding amounts of vehicle, Fsk or Fsk+Ins) for 30 min at 37°C. H_2_O_2_ was then added to a final concentration of 1 mM for exactly 1 min at room temperature with gentle agitation. The reaction was stopped by washing cells three times with PBS containing 5 mM Trolox, 10 mM sodium ascorbate and 10 mM sodium azide. The negative controls which are cells expressing PLIN1-APEX2-V5 were treated identically, except for the 1 min H_2_O_2_ addition step.

### Immunofluorescence staining

After APEX2 labeling, cells were washed with PBS once, fixed with 4% paraformaldehyde in PBS for 15 min at room temperature and washed with PBS three times. Cells were permeabilized and blocked in PBS containing 1% BSA and 0.3% Triton X-100 for 30 min at room temperature and incubated with primary antibody in permeabilization and blocking buffer overnight at 4°C. Cells were washed with PBS three times. Secondary antibodies conjugated to Alexa Fluor-568 or Alexa Fluor-647 were used at 1:1000 dilution for 1 hour at room temperature. To stain lipid droplets, cells were incubated with LipidTOX Deep Red Neutral Lipid Stain or LipidTOX Green Neutral Lipid Stain (1:200 dilution) in PBS for 30 min. For nuclear staining Hoechst 33342 was used in permeabilization and blocking buffer for 5 min. Coverslips were mounted using Prolong Gold Antifade Mountant and imaged using Zeiss LSM 800 or Zeiss Axiovert 200M Florescence microscope. Images were analyzed using ImageJ (FIJI).

### Immunoblotting

After APEX2 labeling, cells were washed with PBS twice at room temperature, and 1 ml of boiling 2% SDS solution supplemented with 1X Halt protease and phosphatase inhibitor cocktail was added to each 10-cm dish. Cell lysates were scrapped into 5-ml Eppendorf tubes and sonicated for 45 sec with 15 sec on/off cycles at 60% amplitude on ice. Cell lysates were incubated at 37°C for 20 min, followed by another incubation at 65°C for 5 min. Protein concentrations of cell lysates were determined using BCA assay. 40 µg of whole cell lysates were combined with Laemmli buffer, heated at 95°C for 10 min and separated on SDS-PAGE gels. Proteins were transferred to PVDF membranes. Membranes were blocked in PBST containing 5% (w/v) skim milk for 1 hour at room temperature and incubated with primary antibodies in PBST containing 5% BSA at 4°C overnight. Primary antibodies were detected with HRP-conjugated secondary antibodies diluted in PBST containing 5% BSA at room temperature for 45 min. For identification of biotinylated proteins, membranes were incubated with Neutravidin-HRP in PBST containing 5% BSA at room temperature for 30 min. HRP activity was visualized after incubation of SuperSignal West Pico PLUS substrate and imaging on Bio-Rad ChemiDoc imaging system. The following antibodies and reagents were used: anti-V5 tag (Invitrogen, 1:1000 dilution), anti-PLIN1 (Cell Signaling Technology, 1:1000 dilution), anti-Phospho-PLIN1-Serine 522 (Vala Sciences, 1:4000 dilution), anti-GAPDH (Cell Signaling Technology, 1:1000), anti-HSL (Cell Signaling Technology, 1:1000 dilution), anti-ATGL (Cell Signaling Technology, 1:250 dilution), Pierce High Sensitivity Neutravidin-HRP (Thermo Fisher, 1:5000 dilution), goat anti-rabbit HRP conjugated secondary antibody (Invitrogen, 1:2500 dilution for HSL and ATGL blots, 1:10000 dilution for other immunoblots), goat anti-mouse HRP conjugated secondary antibody (Invitrogen, 1:10000).

### Glycerol assay

Cell culture media were collected at specific time points and glycerol in media was measured using free glycerol reagent (Sigma-Aldrich) according to the manufacturer’s protocol.

### Enrichment of biotinylated proteins

To enrich biotinylated proteins, 1800 µg of protein samples in 2% SDS solution were diluted 1:10 with RIPA lysis butter without SDS (50 mM Tris-HCl, pH 7.5, 150 mM NaCl, 0.5% sodium deoxycholate, 1% TritonX-100 in distilled water). 200 µl of Cytiva SpeedBeads magnetic neutravidin coated particles were washed 2X with RIPA lysis buffer (50 mM Tris-HCl, pH 7.5, 150 mM NaCl, 0.1% SDS, 0.5% sodium deoxycholate, 1% TritonX-100 in distilled water) and added to the diluted protein samples, followed by overnight incubation at 4°C with rotating. Next day, beads were washed 4X with RIPA buffer to remove nonspecific binding and were resuspended in 170 µl of RIPA buffer. 150 µl of beads suspensions were sent to the Proteomics and Mass Spectrometry Facility at University of Georgia for LC-MS/MS analysis. 20 µl of beads suspensions were used for elution of biotinylated proteins in 60 µl of 3X Laemmli buffer containing 2 mM biotin and 20 mM DTT with heating at 95°C for 10 min. 30 µl of eluates were loaded on SDS-PAGE gels for western blot analysis using Neutravidin-HRP or antibodies indicated in each figure.

### On-beads digestion and LC-MS/MS analysis of biotinylated proteins

Beads suspensions were sent to Proteomics and Mass Spectrometry Facility at University of Georgia for proteomic analysis. The beads were washed with 200 µl of 20 mM ammonium bicarbonate, vortexed, and centrifuged at 1000 x g for 2 minutes. The solvent was removed and replaced with 20 mM ammonium bicarbonate. This wash step was repeated for another 5 times. The proteins on the beads were then digested with 0.2 µg of sequencing grade trypsin in about 40 µl of bicarbonate buffer overnight at room temperature. Next day, 100 µl of water was added to quench trypsin digestion and then the tryptic peptides in the supernatant were collected. The peptides were dried down in a vacufuge and resuspended in 10 µl of 2% acetonitrile containing 0.1% formic acid for LC-MS/MS.

Samples were analyzed on a Thermo-Fisher LTQ Orbitrap Elite Mass Spectrometer coupled with a Proxeon EASY-nLC system. Briefly, 1 µl of enzymatic peptides were loaded into a reversed-phase column (self-packed 100 µm ID column with 200 Å 5 µM Bruker MagicAQ C18 resin, ~15 cm long), then directly eluted into the mass spectrometer at a flow rate of 450 nl/min. The two-buffer gradient elution (0.1% formic acid as buffer A and 99.9% acetonitrile with 0.1% formic acid as buffer B) starts with 2% B, holds at 0%B for 2 minutes, then increases to 30% B in 60 minutes, to 50% B in 10 minutes and to 95% B in 10 minutes. The data-dependent acquisition (DDA) method was used to acquire MS data. A survey MS scan was acquired first, and then the top 10 ions in the MS scan were selected for following CID MS/MS analysis. Both MS and MS/MS scans were acquired by Orbitrap at the resolutions of 120,000 and 15,000, respectively.

### Mass spectrometry data analysis by MaxQuant

For label-free quantification, MaxQuant(40) algorithm (version 1.6.5.0) was used for the identification and quantification of proteins and peptides. The raw data files were searched against the UniProt human proteome database (protein count: 78,120, Proteome ID: UP000005640) with an additional laboratory contaminant provided by MaxQuant. Carbamidomethyl cysteine was searched as a fixed modification. N-terminal protein acetylation and oxidized methionine were searched as variable modifications. The trypsin specificity was set to allow cleavages N-terminal to proline and a maximum of two missed cleavages. First search peptide tolerance was 20 ppm, and for the main search, 6 ppm. The FTMS MS/MS match tolerance was 20 ppm and the top MS/MS peaks per 100 Da was set to 12. The minimum peptide length was set to 7. The peptide spectrum match (PSM) and protein false discovery rates were set to 1%. Protein quantification was based on unmodified, N-terminally acetylated peptides and peptides with oxidized methionine.

### Statistical analysis of label-free proteomics data using ProVision

Downstream statistical analysis was performed using ProVision(41) (https://provision.shinyapps.io/provision/) using ProteinGroup.txt file generated by MaxQuant. The method details were as follows: The contents within column named “Fasta headers” were deleted and the contents within column named “Majority protein IDs” were replaced with the contents within column named “Gene names” in ProteinGroup.txt before it was uploaded to the ProVision. LFQ intensity was chosen for quantification. Proteins that been identified only by a single peptide were filtered out. The LFQ data was converted to Log_2_ scale and subtracted by the median. Samples were grouped by conditions and the proteins with at least 2 values per replicate in at least one group were retained for later statistical analysis. Missing values were imputed using the ‘Missing not At Random’ (MNAR) method, which uses random draws from a left-shifted Gaussian distribution of 1.8 standard deviation apart with a width of 0.3. The limma package from R Bioconductor was used to generate a list of differentially expressed proteins for each pair-wise comparison. A cutoff of the adjusted p-value of 0.05 (Benjamini-Hochberg method) along with a |log_2_ fold change| of 1 has been applied to determine significantly regulated proteins in each pairwise comparison.

### Analysis of LD proteome

Heatmap was generated using the interactive tool Morpheus (Broad Institute). The matrix containing the imputed LFQ intensity values of identified proteins was used as input data for Pearson correlation analysis to compute the nearest neighbors of PLIN1. Area-proportional Venn diagram was generated using the lists containing all the identified proteins or high confidence LD proteins in human primary adipocytes or human cancer cell lines (U2OS and Huh7) using BioVenn (42). KEGG pathway and Gene Ontology (GO) analysis of high confidence LD proteomes from human primary adipocytes and human cancer cell lines were performed and visualized using ShinyGO 0.77 (http://bioinformatics.sdstate.edu/go/)

### siRNA-mediated knockdown of 14-3-3 proteins

For silencing of 14-3-3β, three Silencer^®^ Select siRNA oligos specifically targeting YWHAB (Thermo Fisher, siRNA ID: s14961, s14962, s14963) were used. On day 6 of differentiation, transfections were performed using Endo-Porter (Gene Tools). The final concentrations of siRNA and Endo-Porter were 20 nM and 7.5 µM, respectively. 48 hrs after transfection, cells were stimulated with Fsk without or with the presence of insulin and then harvested for RT-qPCR or western blot analysis. In addition, culture media were collected 6hrs or 24 hrs post stimulation for glycerol assay. The sequences of siRNAs targeting the four isoforms from 14-3-3 family used in this study are listed in the following tables.

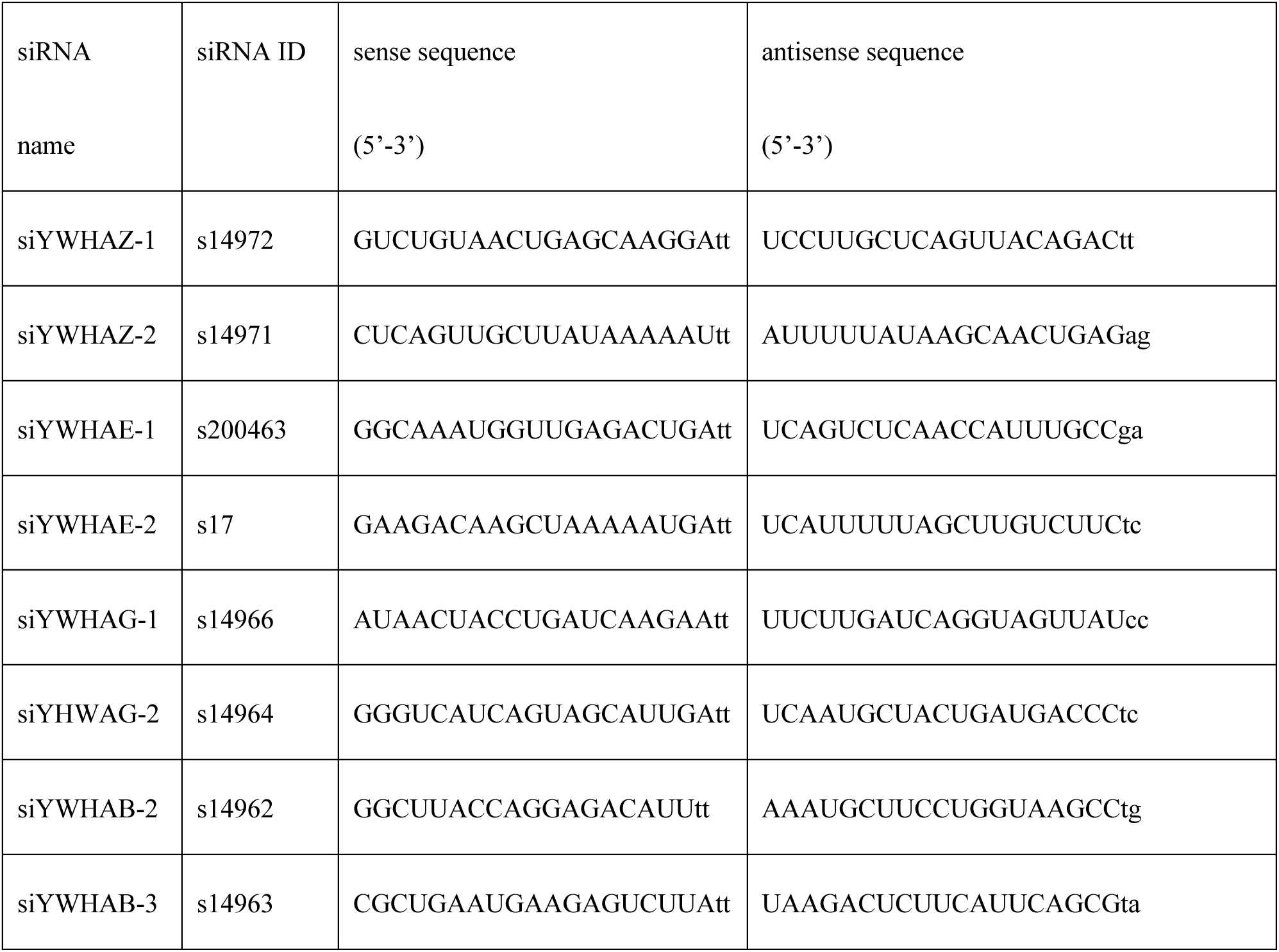

### RT-qPCR analysis

Total RNA was purified using Direct-zol RNA Miniprep Kit (Zymo Research) following the instructions of the manufacturer or using the TriPure Isolation Reagent. Reverse transcriptions were performed for 1 µg RNA using iScript cDNA Synthesis Kit (Bio-Rad) according to the manufacturer’s protocol. cDNA was loaded in duplicate, and qPCRs were performed with iQ SYBR Green Supermix (Bio-Rad) using CFX96 Touch Deep Well Real-Time PCR Detection System (Bio-Rad). The qPCR primers used in this study are listed in the following table.

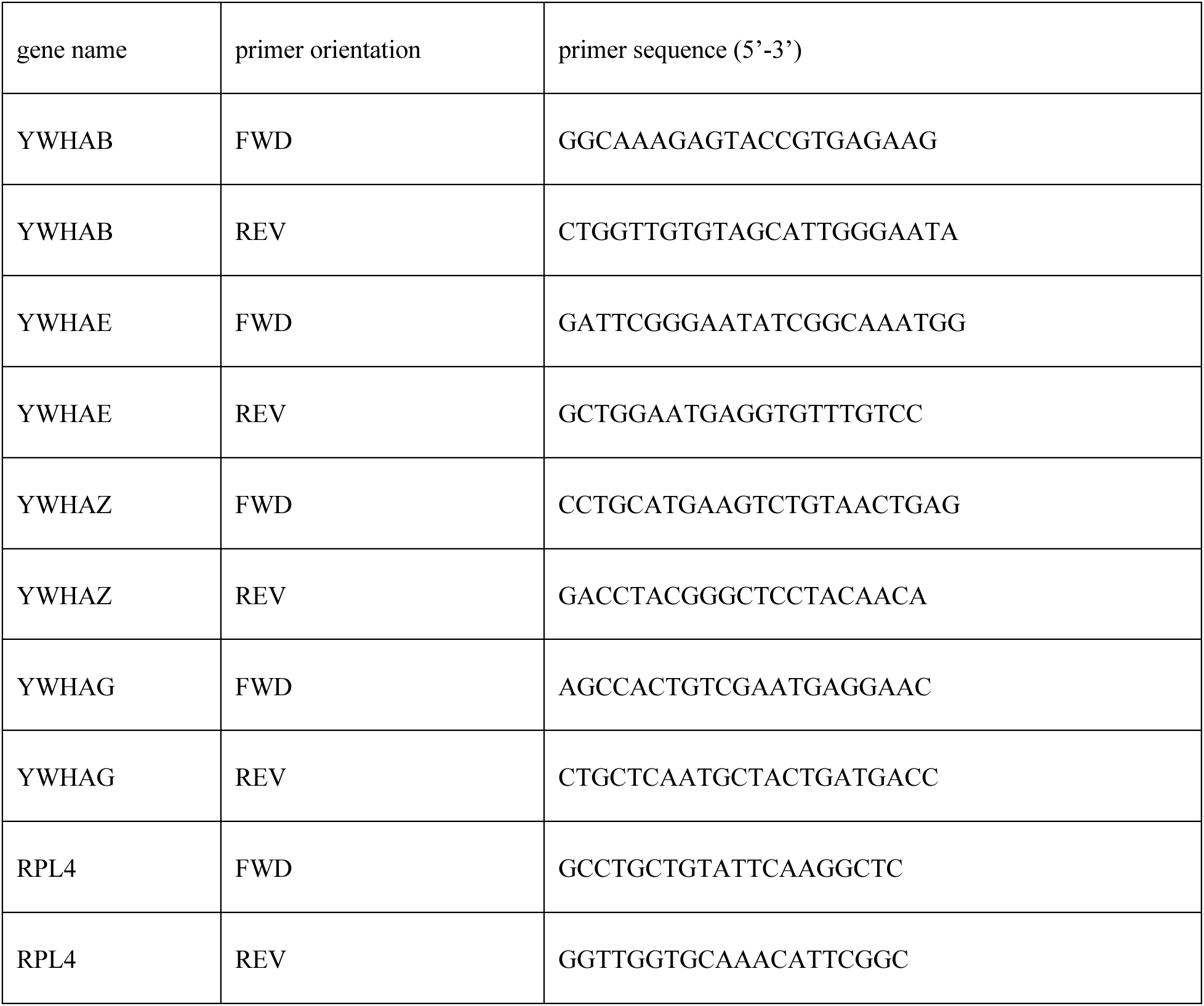
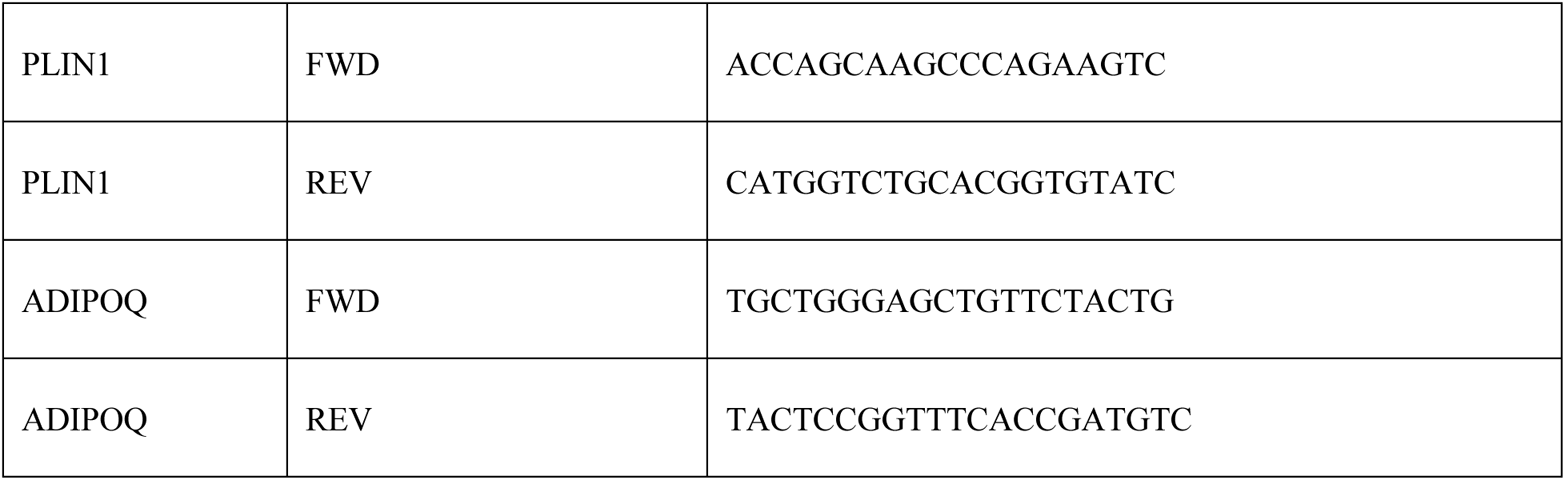

### RNP transfection in progenitors of human primary adipocytes

Neon Transfection System 10 µL Kits (ThermoFisher) were used for ribonucleoprotein (RNP) nucleofection according to the manufacturer’s protocol. Briefly, RNP complexes consisting of 4 µM sgRNA (IDT or Synthego) and 3 µM SpyCas9 protein (PNA Bio) were prepared in Buffer R provided by the Neon Transfection System Kit to a volume of 6 µl. Progenitor cells were trypsinzed, washed and resuspended in Buffer R to a density of 2×10^5^ cells per 6 µl. Equal volume of RNP mix and cell suspension were mixed using 10 µl Neon tip and electroporated using the optimized parameters (voltage: 1350V, width of pulse: 30 ms, number of pulse: 1). After electroporation, the cells were plated immediately in 12-WP containing 1 ml of complete media, grown to confluence, and differentiated into adipocytes according to the methods mentioned in <u>cell culture from human adipose tissue explants</u> for downstream experiments. The sequences of sgRNAs used in this study are listed in the following table.

sgRNA targeting YWHAB

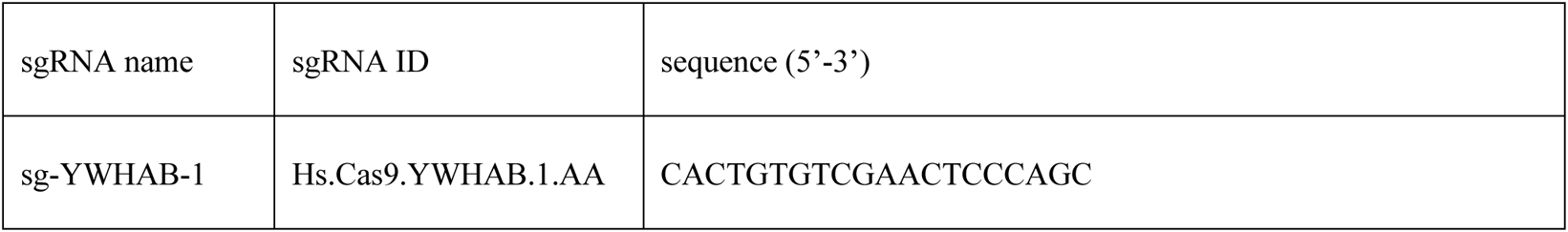
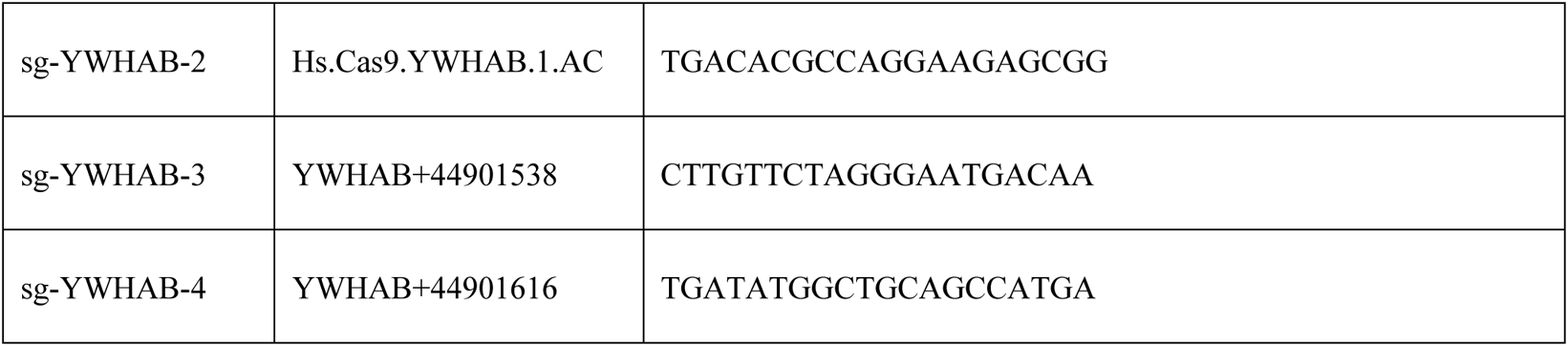

### Indel analysis by TIDE

The electroporated progenitors were subjected to adipogenic differentiation for 8 days and the genomic DNA was isolated using QuickExtract DNA Extraction Solution (Lucigen). PCR reactions were conducted with 50 ng of genomic DNA and following primer pairs spanning the sgRNA target site in KAPA HiFi HotStart ReadyMix (Roche) according to the manufacturer’s instructions. The PCR products were purified by QIAquick PCR Purification Kit and summitted Genewiz for Sanger Sequencing. The control and sample sequence data files were analyzed using TIDE webtool (htEtps://tide.nki.nl/#about) to quantify indel frequencies. The PCR primers designed for sgRNAs targeting YWHAB are listed in the following table.

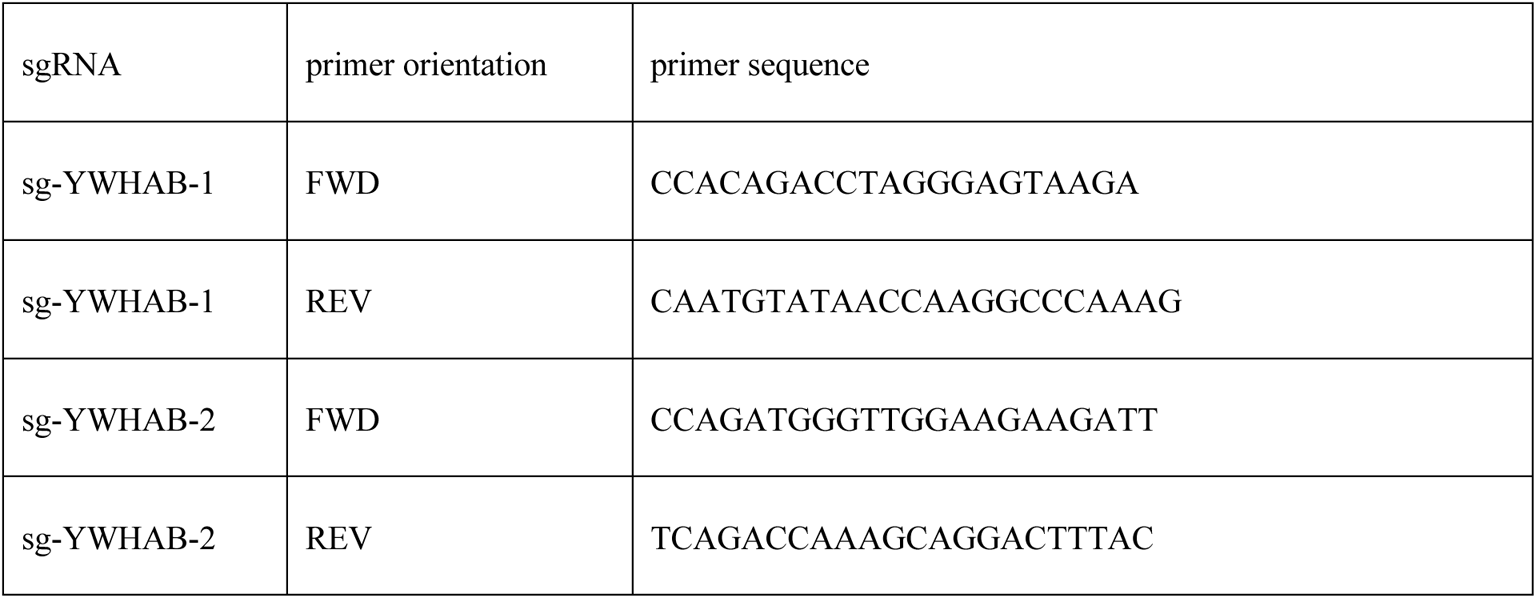
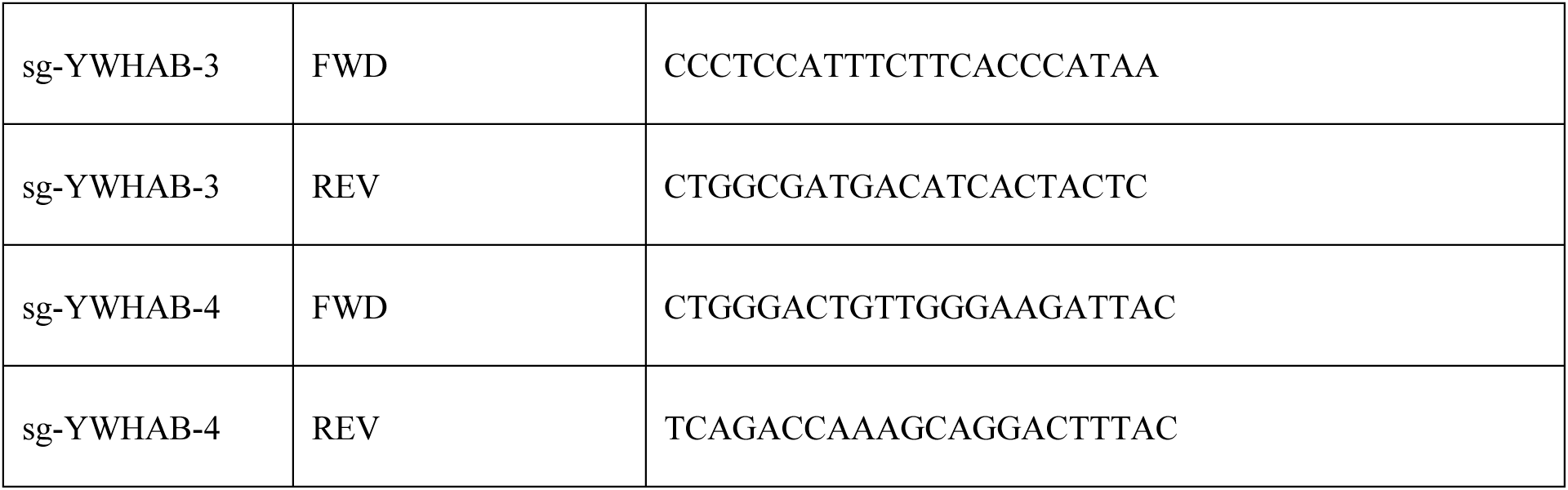

### Cell viability assays

CellTiter-Glo 2.0 cell viability assay (Promega, Cat# G9243) was performed according to the manufacturer’s protocol. The fluorescence and luminescence signals were measured using a Safire 2 microplate reader (Tecan).

### In situ proximity ligation assay

In situ proximity ligation assays (PLAs) were performed using Duolink in situ red starter kit according to the manufacturer’s instructions (Millipore Sigma, DUO92101). Briefly, the electroporated cells were subjected to differentiation for 8 days. Cells were then fixed in 4% paraformaldehyde in PBS for 15 min at room temperature (RT) and washed with PBS three times. Cell were permeabilized with 0.3% Triton X-100 in for 30 min at RT. Cells were blocked by Duolink in situ blocking solution for 1 h at 37 °C. Then, cells were incubated with primary antibodies: anti-PLIN1 (1:200 dilution, Cell Signaling Technology, #9349) and anti-YWHAB (1:100 dilution, Santa Cruz Biotechnology, sc-25276) for 2 h at 37 °C and washed twice with 1X Duolink wash buffer A. Cells were incubated with anti-rabbit (PLUS) and anti-mouse (MINUS) probes for 1 h at 37 °C and then washed twice with 1X Duolink wash buffer A. Ligation solution was added to each sample for 30 min at 37 °C and the samples were washed twice with 1X Duolink wash buffer A. Amplification solution was added for 1 h 40 min at 37 °C and the samples were washed with 1X Duolink wash buffer B and 0.01X Duolink wash buffer B. The samples were mounted with Duolink In Situ Mounting Media with DAPI. The images were taken under ZEISS Axiovert 200M Fluorescence microscope. Quantification was performed using ImageJ (FIJI) software.

### Immunoprecipitation

Lenti-X 293T cells were transfected with pCDNA3.1 plasmids expressing HA-tagged YWHA isoforms using Lipofectamine 3000 (Thermo Fisher Scientific) according to the manufacturer’s protocol and cultured for 48 h before lysis in 2% SDS buffer. The cell lysates in 2% SDS were diluted with RIPA lysis butter without SDS (50 mM Tris-HCl, pH 7.5, 150 mM NaCl, 0.5% sodium deoxycholate, 1% TritonX-100 in distilled water) to a final concentration of 0.2% SDS and sonicated. Protein concentrations of cell lysates were determined using BCA assay. A total of 1mg protein lysate was incubated with 50 µl suspension of monoclonal anti-HA agarose (Millipore). After overnight incubation, the samples were washed with RIPA lysis buffer four times and the proteins bound to the agarose were eluted with 25 µl of 2X Laemmli buffer by boiling at 95 °C for 3 min. 5 µl of eluates were loaded on SDS-PAGE gels for western blot analysis using the following antibodies: anti-HA Tag (Cell Signaling Technology, #3724), anti-YWHA (pan, Cell Signaling Technology, #8312), YWHAB (Abcam, ab15260), YHWAB (Santa Cruz Biotechnology, sc-25276), YWHAE (Cell Signaling Technology, #9635), YWHAE (Santa Cruz Biotechnology, sc-23957), YWHAZ (Santa Cruz Biotechnology, sc-293415), YWHAZ (Abcam, ab51129), YWHAG (Cell Signaling Technology, #5522S), YWHAG (Novus Biologicals, NB100-407SS), and YWHAG (Santa Cruz Biotechnology, sc-398423).

## ACKNOWLEDGEMENTS

We thank Chau-Wen Chou from the Proteomics and Mass Spectrometry Core Facility at the University of Georgia for providing the LC-MS/MS service and raw files. This study was supported by NIH grants DK089101 and DK123028 to SC.

---

**Figure S1.**
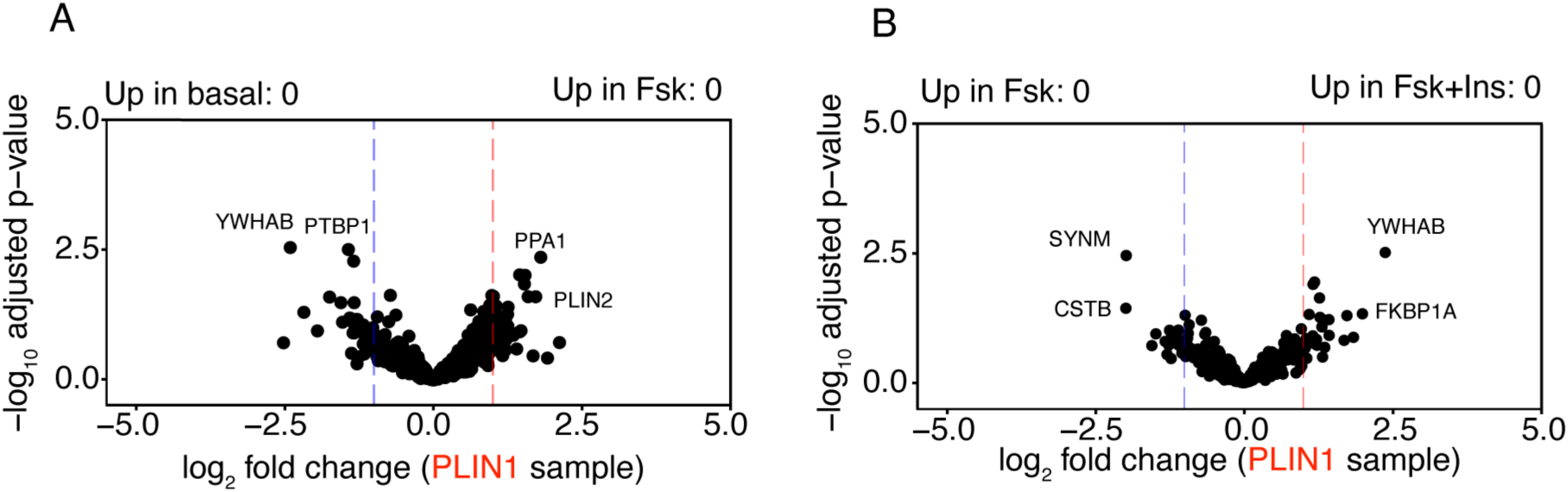
Analysis of proteins biotinylated by PLIN1-APEX2 under different lipolytic conditions. Volcano plots of biotinylated proteins enriched by PLIN1-APEX2 under basal or Fsk-stimulated conditions (A), or under Fsk or Fsk+Ins conditions (B). No proteins were identified using Benjamini-Hochberg-adjusted p-value < 0.05 and log2 fold change ≥ 1.0.

**Figure S2.**
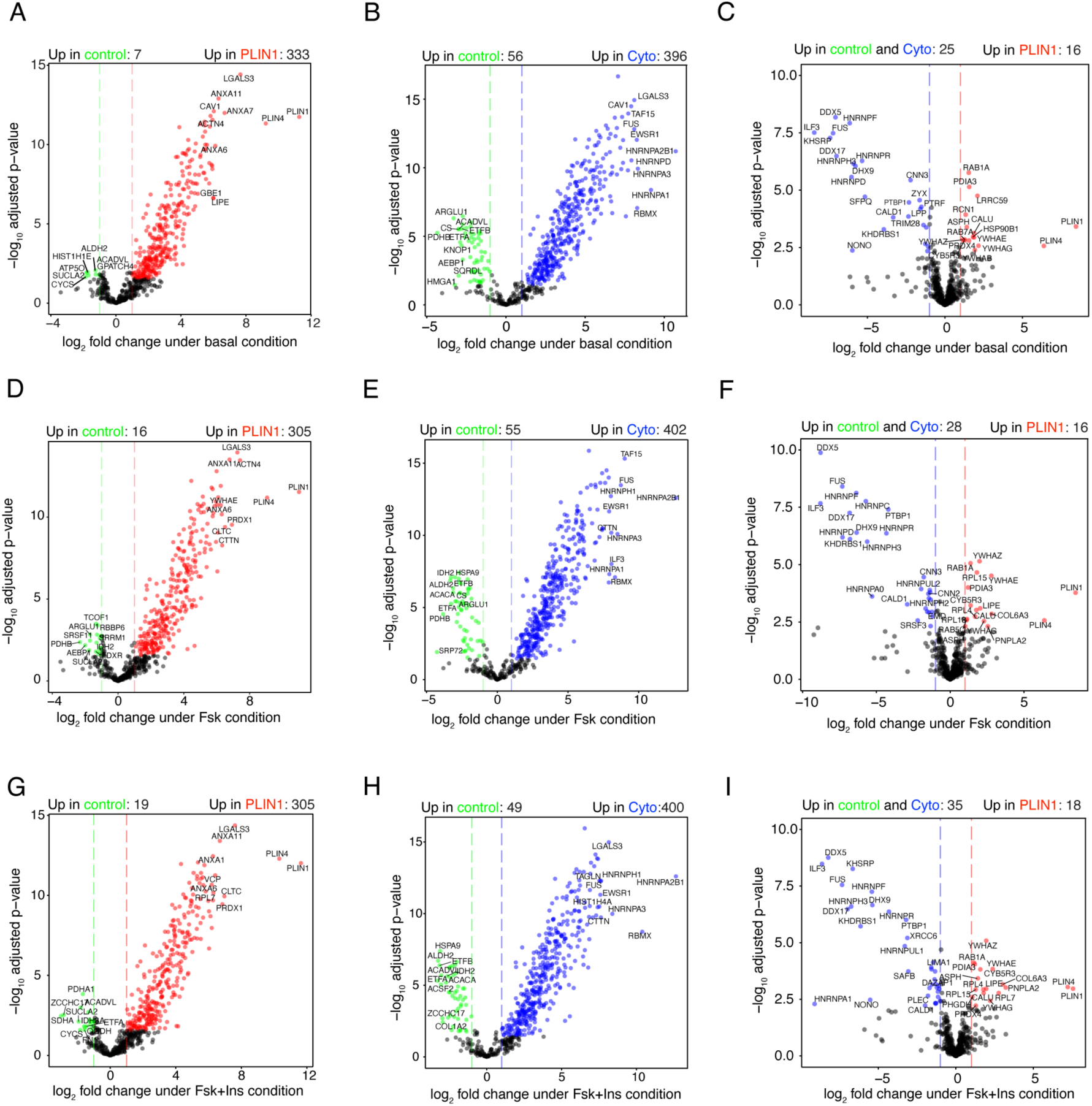
Differential enrichment of biotinylated proteins under different lipolytic conditions. Volcano plots of biotinylated proteins enriched by PLIN1-APEX2 (A,D,G) or Cyto-APEX2 (B,E,H) compared to negative control samples, or by PLIN1-APEX2 compared to combined negative control and Cyto-APEX2 (C,F,I), under basal (A-C), Fsk (D-F), or Fsk+Ins (G-I) treatment conditions.

**Figure S3.**
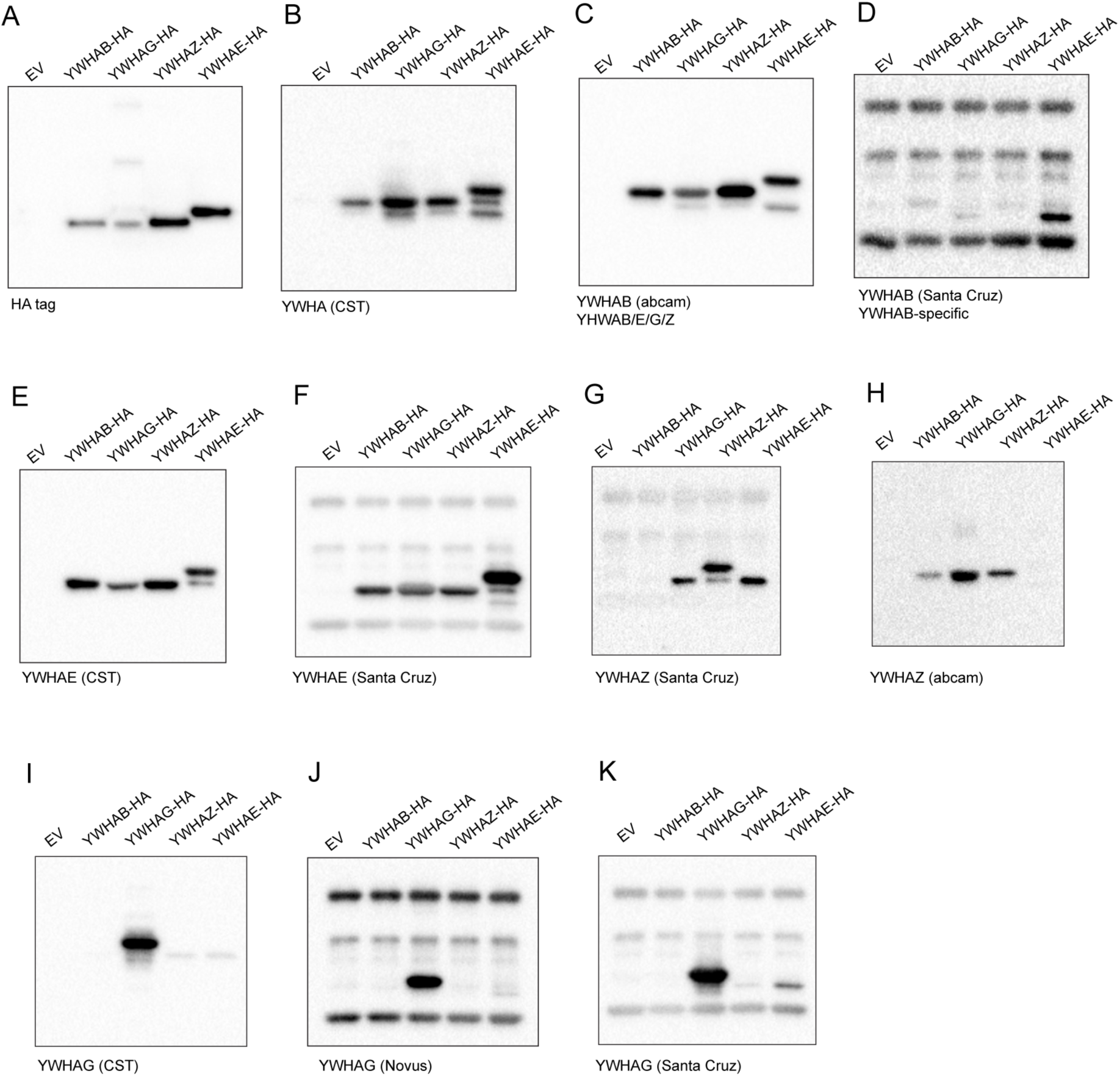
Specificity of commercial antibodies targeting 14-3-3 protein isoforms. 293T cells were transfected by pcDNA3.1 empty vector or pcDNA3.1 vector containing YWHAB-HA, YWHAG-HA, YWHAZ-HA, or YWHAE-HA. Immunoprecipitated HA-tagged proteins were detected by western blotting using antibodies against: (A) HA tag, (B) YWHA pan from abcam, (C) YWHAB from abcam, (D) YWHAB from Santa Cruz, (E) YWHAE from CST, (F) YWHAE from Santa Cruz, (G) YWHAZ from abcam, (H) YWHAZ from Santa Cruz, (I) YWHAG from CST, (J) YWHAG from Novus, and (K) YWHAG from Santa Cruz. The heavy chain and light chains of mouse anti-HA antibody used for immunoprecipitation were also detected when primary antibodies to 14-3-3 isoforms were produced in mouse (D,F,G,J,K).

